# MODEL PREDICTIVE CONTROL OF CANCER CELLULAR DYNAMICS: A NEW STRATEGY FOR THERAPY DESIGN

**DOI:** 10.1101/2022.03.14.484268

**Authors:** Benjamin Smart, Irene de Cesare, Ludovic Renson, Lucia Marucci

**Affiliations:** Department of Mechanical Engineering, Imperial College London, London SW7 1AY, U.K.; Department of Engineering Mathematics, University of Bristol, Woodland Road, Bristol BS8 1UB, U.K.; BrisSynBio, Life Sciences Building, University of Bristol, Tyndall Avenue, Bristol BS8 1TQ, U.K.; School of Cellular and Molecular Medicine, University of Bristol, University Walk, Bristol BS8 1TD, U.K.

**Keywords:** Adaptive Model Predictive Control (MPC), Combination Therapies, Cybergenetics, External Feedback Control, Non-Small Cell Lung Cancer (NSCLC)

## Abstract

Recent advancements in Cybergenetics have led to the development of new computational and experimental platforms that enable to robustly steer cellular dynamics by applying external feedback control. Such technologies have never been applied to regulate intracellular dynamics of cancer cells. Here, we show *in silico* that adaptive model predictive control (MPC) can effectively be used to steer signalling dynamics in Non-Small Cell Lung Cancer (NSCLC) cells to resemble those of wild-type cells, and to support the design of combination therapies. Our optimisation-based control algorithm enables tailoring the cost function to force the controller to alternate different drugs and/or reduce drug exposure, minimising both drug-induced toxicity and resistance to treatment. Our results pave the way for new cybergenetics experiments in cancer cells for drug combination therapy design.

## 1 Introduction

Cybergenetics is a recent field of synthetic biology, which refers to the forward engineering of complex phenotypes in living cells applying principles and techniques from control engineering [1].

Three main approaches have been proven to be effective for the control of different processes (such as gene expression and cell proliferation), namely: i) open- or closed-loop controllers embedded into cells by means of synthetic gene networks [2–6]; ii) external controllers, where the controlled processes are within cells, while the controller (either at single cell or cell-population level) and the actuation functions are implemented externally via microfluidics-optogenetics/microscopy-flow cytometry platforms and adequate algorithms for online cell output quantification and control [7–16]; iii) multicellular control, where both the control and actuation functions are embedded into cellular consortia [17–21]. Plenty of examples of embedded controllers have been engineered across different cellular chassis; instead, applications of external and multicellular controllers in mammalian cells are scarce and either just theoretical or limited to proof of concepts.

Here, we propose to apply cybergenetics, in particular external feedback control, to predict combinations of drugs (i.e. control inputs) which can bring dysregulated cellular variables (i.e. gene expression, control output of the system) within tightly controlled ranges in cancer cells. We take Non-Small Cell Lung Cancer (NSCLC) as an example; using a previously proposed differential equations mathematical model describing the dynamics of the EGFR and IGF1R pathways, we show *in silico* that external feedback controllers can effectively steer intracellular gene expression dynamics in cancer cells to resemble those of wild-type cells.

The use of feedback control is advantageous as it enables coping with changes in both steady-state levels and in the temporal dynamics of genes involved in dysregulated signalling cascades. The control action is implemented by means of adaptive model predictive control (MPC), thus not requiring an exact model of the system; this is particularly advantageous in biological applications, where the derivation of detailed models can be time-consuming and troublesome [22–24].

The possibility to control single/multiple outputs with one or more inputs can support the design of combination therapies which target different nodes in signalling cascades; this approach can be advantageous to maximise the efficacy of cancer therapies [25]. In this regard, our optimisation-based control algorithm enables tailoring the cost function to force the controller to alternate different drugs and/or reduce drug exposure. The controller should also be able to cope with the crosstalk in signalling pathways, which might be further mechanisms causing drug resistance [26].

In what follows, we demonstrate that adaptive MPC can be used to effectively steer the concentration of several proteins within different signalling pathways of a NSCLC cell in order to tune gene expression, whilst reducing the dose of each drug provided as an input. Our results pave the way for using cybergenetics for cancer treatment design.

## 2 Methods

### 2.1 Control Scheme Used in External Feedback

We applied a feedback controller to regulate the concentrations of two downstream genes (ERK and Akt) of the mTOR and MAPK pathways as modelled in [27], and as shown in Figure 1.

**Figure 1:**
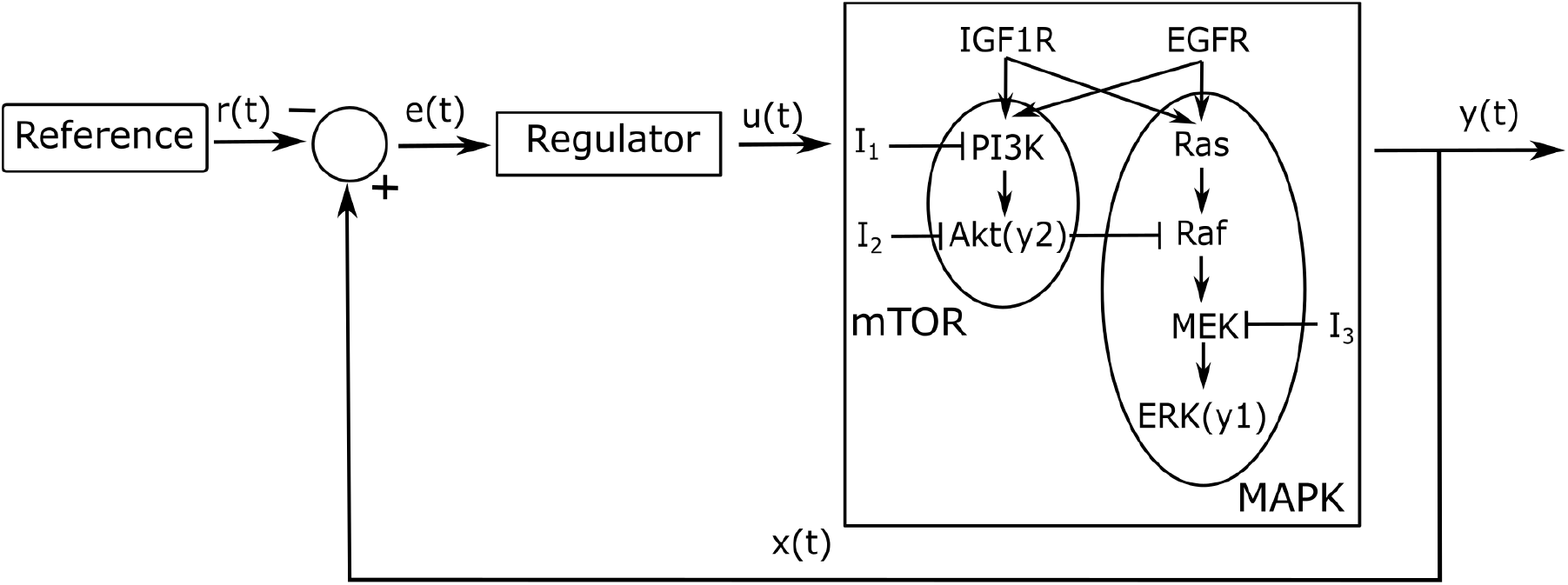
A control scheme including three inputs (*I*_1_, *I*_2_, *I*_3_) that interact with the mTOR and MAPK pathways. Two observable protein concentrations, y1(ERK) and y2(Akt), are used as control outputs for the two pathways. The regulator used throughout this project is an adaptive MPC program which attempts to steer the concentrations of the outputs to the transient response of the wild type cell, set as the control reference, as shown in Figure 2.

Figure 2 shows the response of y1(ERK) and y2(Akt) in a wild type ~ and in a NSCLC ~ cell to a phosphorylation of EGFR and IGF1R as modelled by varying the system’s initial conditions as in [27], referred to as an activation. A wild type cells’ activation is modelled using an active concentration of 8000 *μM* for EGFR and 800*μM* for IGF1R, while an activation in NSCLC cells is triggered by an active concentration of 800000 and 400000 *μM* for EGFR and IGF1R, respectively. The term ‘free’ refers to an open loop response (i.e. if no feedback control is applied) in NSCLC cells. It can be seen that y1(ERK) and y2(Akt) activation dynamics are different in cancer vs wild-type cells 2; of note, the activation of y1(ERK) occurs over a timescale of an order of magnitude faster than the activation of y2(Akt).

**Figure 2:**
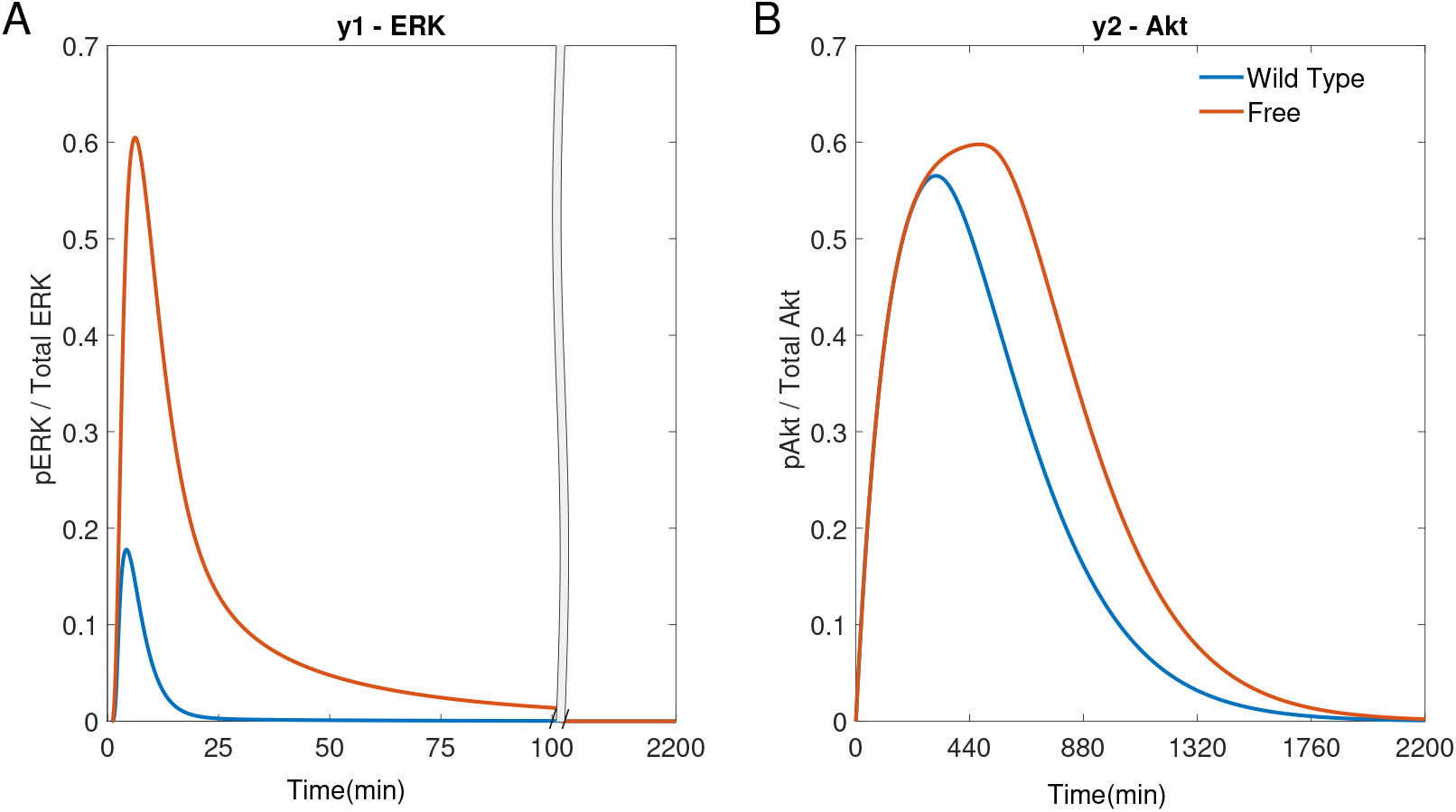
Simulations of the NSCLC model [27]: free NSCLC cell response ~ compared to a wild type cell ~ [27]. (A) The response of y1(ERK). (B) The response of y2(Akt).

The two pathways are both kinase activated cascades, meaning that an activation at the receptors in the cell membrane causes a cascade of phosphorylation in downstream genes. Therefore it is difficult to robustly control the system as, once an error is measured in the outputs, it can be too late to have a significant effect by acting on the internal states higher up the cascade.

A model based controller can interpret a small change in the outputs. On the other hand, model-free feedback strategies with large gains can have undesirable consequences, such as poor dynamic performance. A Proportional controller can be tuned to decrease the control error, but gains have to be carefully chosen, and the user cannot impose desired constrains on the input (Section S8).

Adaptive MPC is used as the model-based control scheme here (Figure 1). The success of MPC relies on the quality of the model used to predict the future behaviour of the system, and on the cost function the controller uses to calculate the optimal inputs to be fed to the process. The novelty of the controller used here lies in the choice of adaptive model and in the cost function used. The MPC implementation is presented in Section S4.

### 2.2 MPC’s Linearised Model

The NSCLC model [27] contains significant non-linear terms and a large number of internal states. An adaptive MPC controller which computes a linear approximation of the NSCLC model at each time step, predicts the future of the internal states and calculates the optimal input profile [28]. The controller then applies the first input of its calculated optimal input profile to the actual system. At the next time step, the controller calculates a new linear model by linearising the NSCLC model. The use of a linear system results in a convex optimisation problem which can be solved quickly. Adaptive MPC is used in all simulations unless stated otherwise.

Alternatively, non-linear MPC could be used; however, it is computationally expensive, and the time needed to compute the next input could be longer than the sampling/actuation temporal intervals (see Section S7).

### 2.3 Improving the Traditional MPC Cost Function

The cost function used by the adaptive MPC algorithm to find the optimal input depends on the current internal state error, **e**_0_, and on the inputs, **u**(*t*). The error, **e**(*t*), is the difference between the reference, **r**(*t*), and the internal states of the NSCLC system, **x**(*t*), as shown in Figure 1. The standard cost function used for linear MPC controllers [29], focuses on how readily the inputs, **u**(*t*), are used and on reducing the proportion error in the states, **e**(*t*).

In order to include both the magnitude and duration of the error, the integral of the state error, ∫ **e**(*t*) *dt*, is added to the standard cost function. It has been shown that it is beneficial to integrate the state error [30], meaning that the controller acts due to these longer, smaller errors in the outputs caused by states higher in the cascade. Moreover, to avoid rapid fluctuations in the control input, **u**(*t*), a differential cost of the inputs is also added to cost function. The derivation of the cost function and a description of the weights of its terms (*α*, *β, γ, η, θ*) can be found in Section S4.

### 2.4 MPC Simulation Parameters

The MPC simulations are reproducible thanks to the deterministic nature of the model and controller, as long as the cost function coefficient weights and other MPC related parameters are kept constant. Table 1 gives a summary of these parameters. Several key parameters are added to the figures’ captions.

**Table 1:**
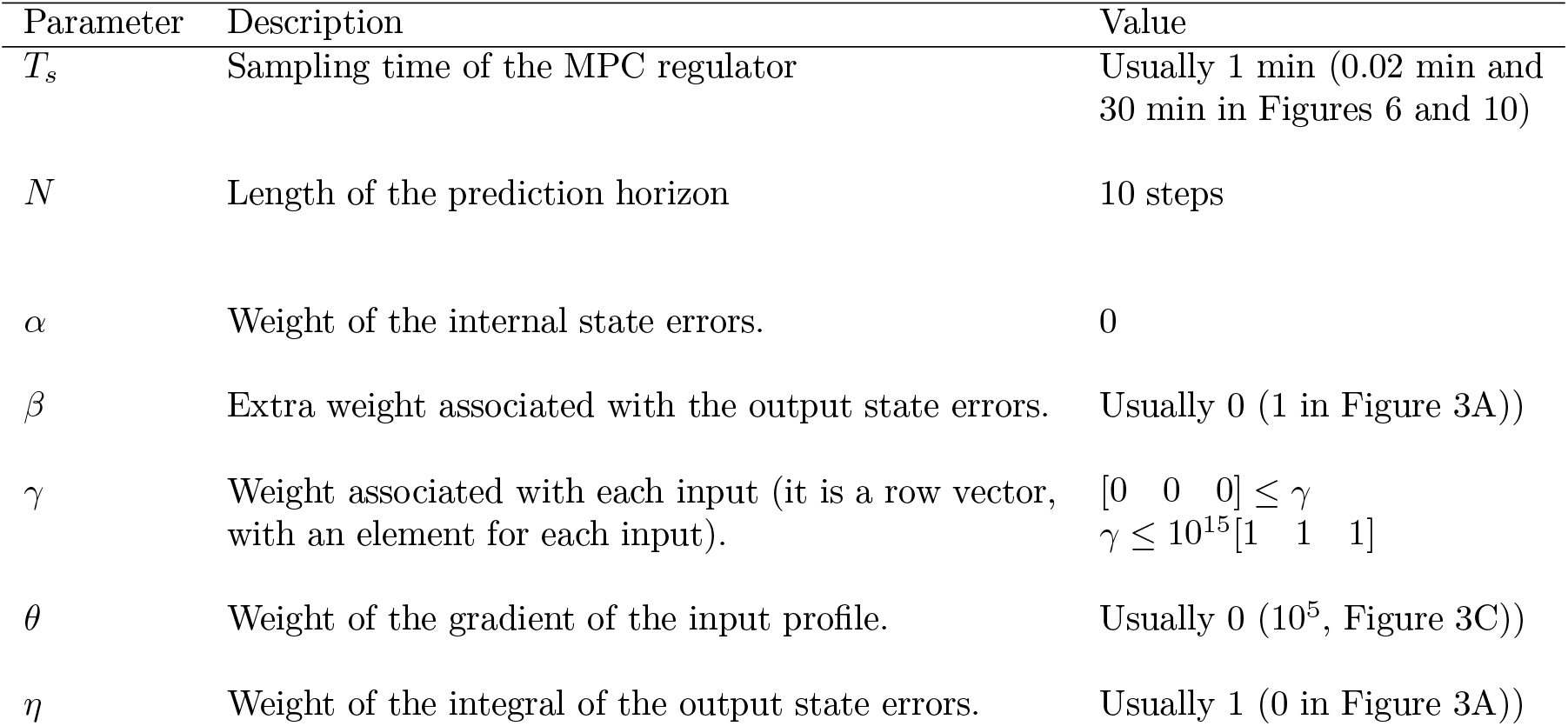
Parameters used for MPC simulations.

### 2.5 Indexes Used to Quantify Control Performance

To assess quantitatively the performance of our controller, we define an Error Index, *EI*. It is the sum of the squared error between the output and the reference for the outputs, as used in [31].

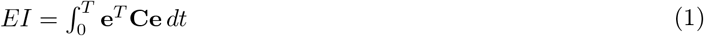

**C** is the output matrix of the linearised NSCLC model. A small *EI* indicates a good performance of the controller.

To quantify the controller effort needed to achieve a certain output, we assess the dose of input drug(s) using a Dose Index, *DI_i_*. It is the integral of the input signal, where **u**(*t*) = [*I*_1_(*t*), *I*_2_(*t*), *I*_3_(*t*)].

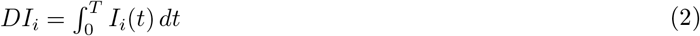

The inputs can never be negative as they are physical concentrations, therefore there is no need to square the input signal.

## 3 Results

Non-Small Cell Lung Cancer (NSCLC) accounts for 80% of lung cancer cases and is characterised by various mutations which usually lead to an overexpression of the EGF and IGF1R receptors. These receptors trigger several cascades including the mTOR and MAPK pathways; their downstream genes FOXO1 and C-FOS regulate cell apoptosis and proliferation. The differential equations-based mathematical model for NSCLC signalling developed in [27] includes the mTOR and MAPK pathways along with some of the reactions between the two pathways, as shown in Figure 1; the model enables comparing gene expression dynamics in wild type vs cancers cell. We chose the downstream genes ERK and Akt (noted as y1 and y2, respectively) as the control outputs for the external feedback loop (Figure 1). The two outputs can be tuned by varying three inputs (*I*_1_, *I*_2_, *I*_3_), which inhibit three specific proteins within the mTOR and MAPK pathways. The pathways can influence each others’ reactions, creating internal feedback loops (crosstalk).

The code used to implement an adaptive MPC program on this NSCLC model used in these simulations is available on GitHub: https://github.com/Ben-Smart/Adaptive_MPC_on_NSCLC.git.

### 3.1 Assessing the Importance of Each Term in the Cost Function

Firstly, Single-Input Single-Output (SISO) simulations were performed. The controller tries to steer the dynamics of either y1(ERK) or y2(Akt) by varying the concentrations of a drug that acts directly on one of the two signalling cascades (*I*_3_ for y1(ERK) and either *I*_1_ or *I*_2_ for y2(Akt)). Figure 3 uses *I*_2_ to regulate y2(Akt), and shows the effect of different cost function terms on the performance of the controller.

**Figure 3:**
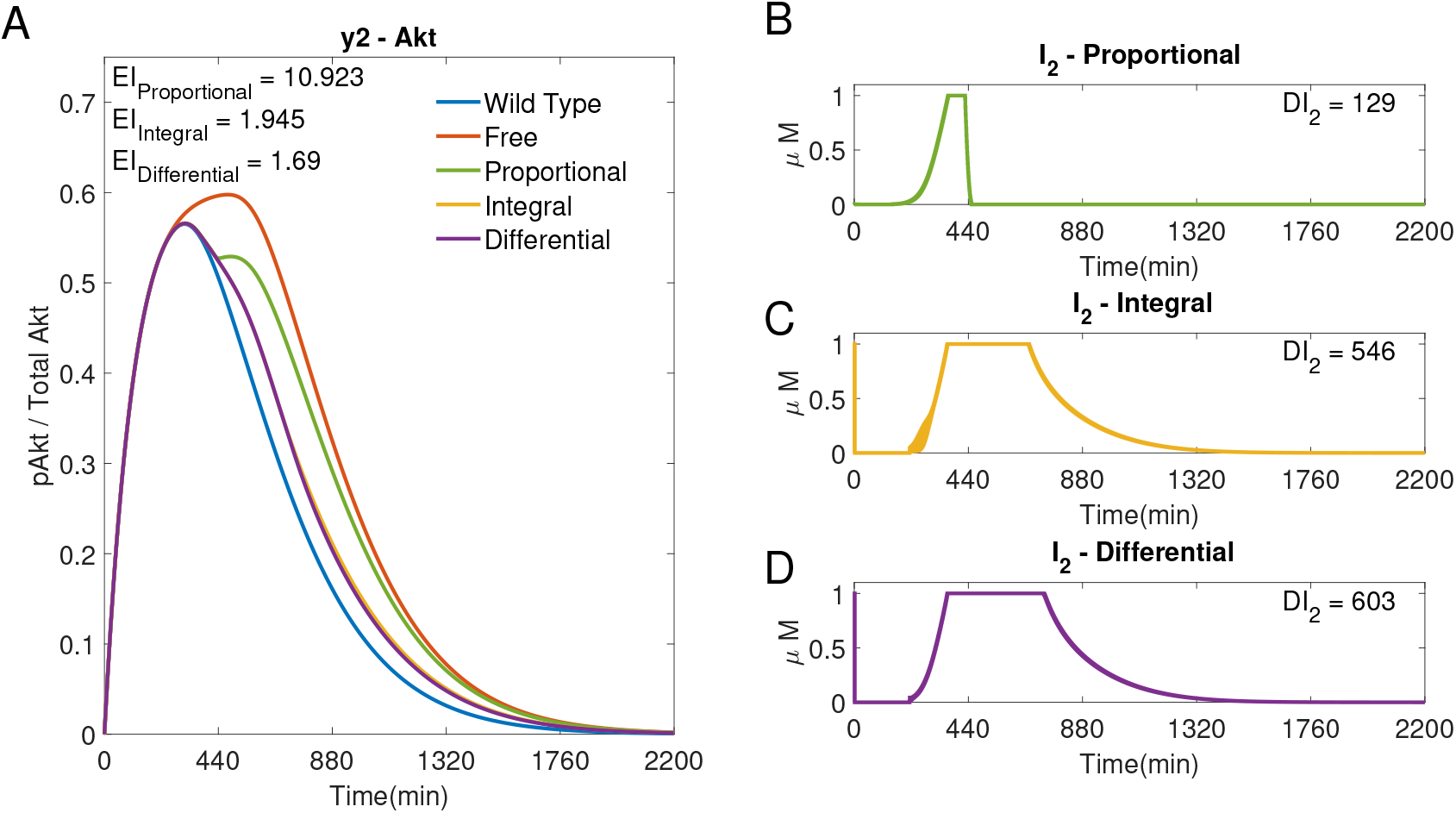
SISO adaptive MPC simulation using *I*_2_ to control y2 (Akt), and comparing different cost functions. (A) The response of y2 to different cost functions. (B) The input profile (*I*_2_) using the proportional error ~ within the cost function. *β* =1, *γ* = [–, 10^5^, –], *θ* = 0 and *η* = 0. (C) The input profile (*I*_2_) using the integral error ~. *β* = 0, *γ* = [–, 10^5^, –], *θ* = 0 and *η* =1. (D) The input profile (*I*_2_) using the integral error and input differential terms ~ of the cost function. *β* = 0, *γ* = [–, 10^5^, –], *θ* = 10^5^ and *η* =1. *T_s_* = 1*min* and *N* =10 for all plots.

Figure 3A shows that using integral terms within the cost function reduces the error in y2(Akt), as compared to the proportional terms (*EI* → 1.945~< 10.923~). However, it can be seen in Figure 3C that the controller using integral terms ~ can cause fluctuations in the input. Such fluctuations are reduced when using a differential cost, which also has a lower Error Index, but higher Dose Index (*EI* → 1.690~< 1.945~< 10.923~, *DI*_2_ → 603~> 546~> 129~).

The derivative term is not included in further simulations; the cost functions used in the following simulations include *β* = 0, *θ* = 0 and *η* = 1, as in Figure 3C, apart from the weight associated with each input, *γ*, that is varied (as indicated in figures’ captions).

#### 3.1.1 Single-Input Single-Output Control with the Chosen Cost Function

Figure 4 shows SISO adaptive MPC simulations for *I*_1_, *I*_2_ and *I*_3_; each drug is used to control the downstream molecule in the cascade it acts on. It can be seen that plots C) and D) of Figure 4 are identical to ∼ in plot A) and plot C) of Figure 3 as these are both SISO responses of *I*_2_ using the chosen cost function (costing the integral and input terms). All of the SISO controllers move the NSCLC response towards the wild type response ~ using a lower dose than just a step of each input at the maximum allowed dose (1*μM*), decreasing the Dose Indexes. The step input response can be found in Section S3. This demonstrates the benefits of using an external feedback loop compared to an open loop response with a static step input.

**Figure 4:**
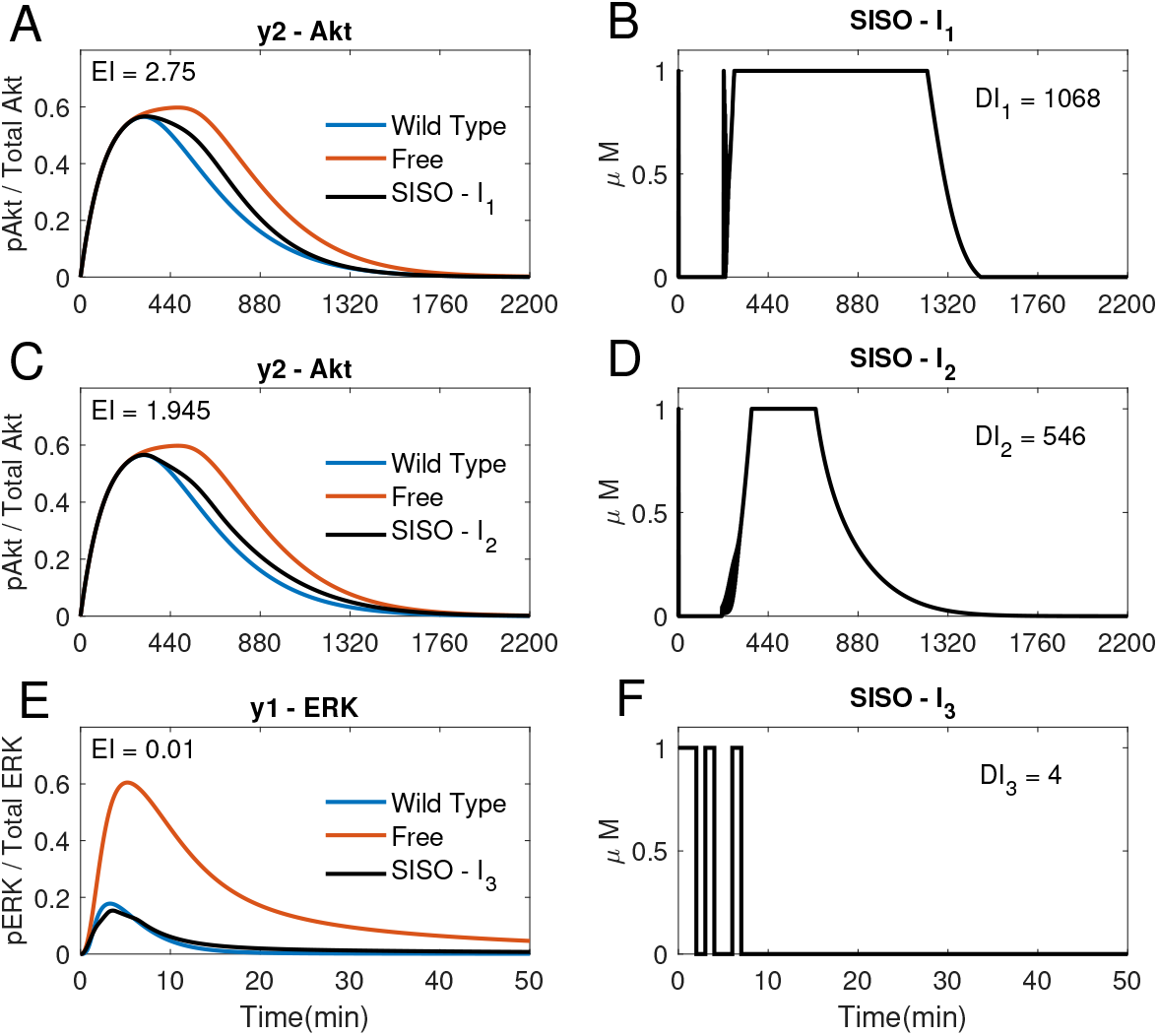
SISO adaptive MPC simulations. (A, B) The response of y2(Akt, in A) to the input of *I*_1_ (shown in B); *γ* = [1, –, –]. (C, D) The response of y2 (Akt, in C) to the input of *I*_2_ (shown in D); *γ* = [–, 10^5^, –]. (E, F) The response of y1(ERK, in E) to the input of *I*_3_ (shown in F); *γ* = [–, –, 10^9^]. (A-F): *T_s_* = 1, N =10, *α* = 0, *β* = 0, *θ* = 0 and *η* =1.

### 3.2 Multi-Input Multi-Output Control

Adaptive MPC can also be used to steer both outputs using all three inputs in Multi-Input Multi-Output (MIMO) simulations, as shown in Figure 5.

**Figure 5:**
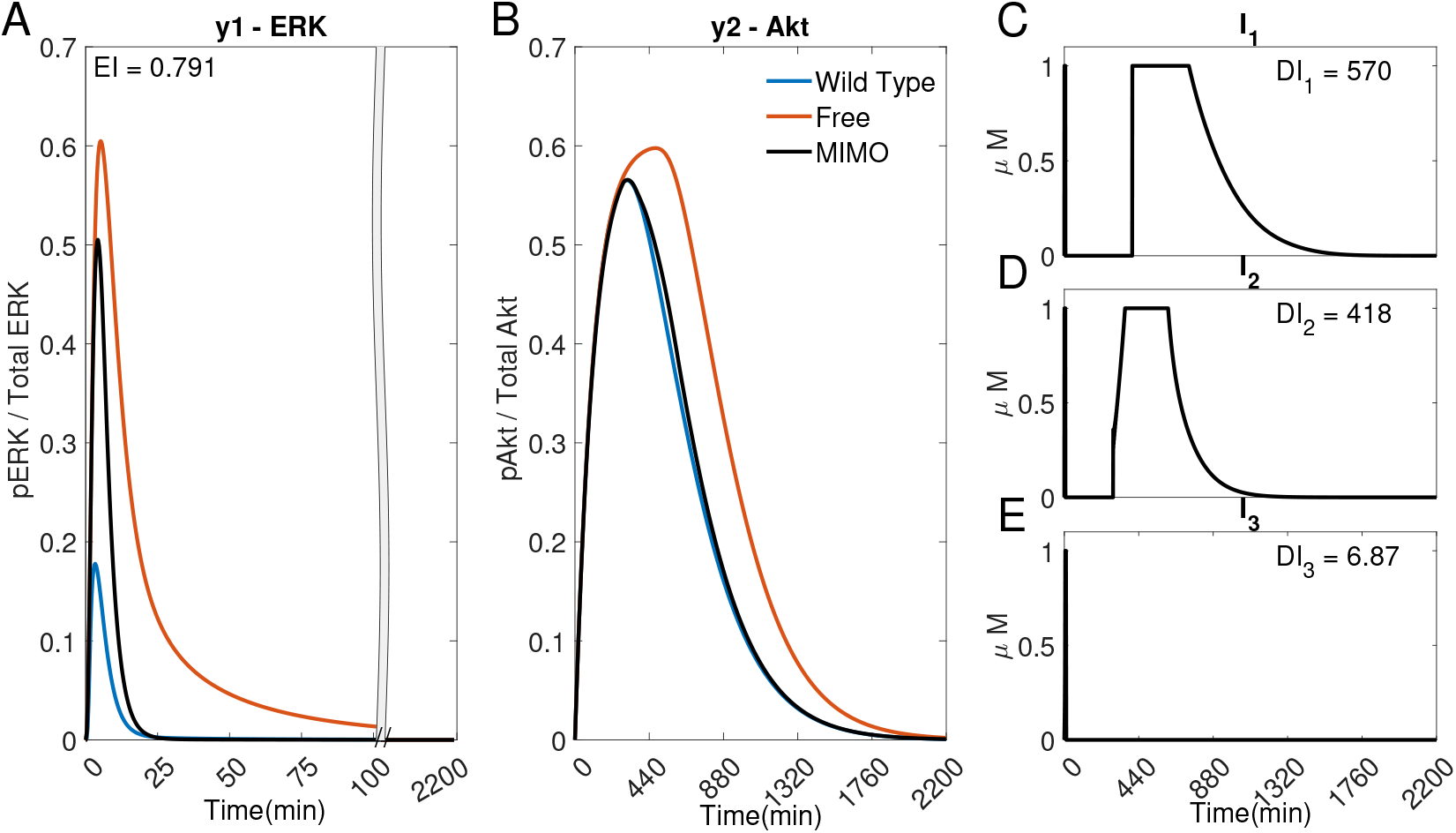
A MIMO adaptive MPC simulation using *I*_1_, *I*_2_ and *I*_3_ to control the concentrations of y1 (ERK) and y2 (Akt). (A) The response of ERK. (B) The response of Akt. (C-E) The inputs used in the simulation. Parameters: *T_s_* = 1*min, N* =10, *α* = 0, *β* = 0, *γ* = [1, 10^5^, 10^9^], *θ* = 0 and *η* =1.

The error of y2 (Akt) (Figure 5B)) is significantly smaller in comparison to Figure 4A and C (*EI* → 0.791 < 1.954 < 2.75), whilst using significantly less *I*_1_ (*DI*_1_ → 570 < 1068) and *I*_2_ (*DI*_2_ → 418 < 546), suggesting that it might be advantageous to use adaptive MPC to predict and apply combination drug profiles.

However, due to the fast dynamics of the MAPK pathway, the output y1(ERK) fails to adequately follow the reference activation curve. Figure 6 shows that if the time step is adequately reduced (for instance, to *T_s_* = 0.02 minutes), the controller can handle the faster dynamics of the pathway and effectively control both outputs (*EI* → 0.001 < 0.791) whilst using a lower dosage of all the inputs (*DI*_1_ → 545 < 570, *DI*_2_ → 297 < 418, *DI*_3_ → 4.35 < 6.87). Figure 6 shows that the controller can overcome the effect of crosstalk, balancing the error in y1(ERk) and the use of *I*_1_ and *I*_2_.

**Figure 6:**
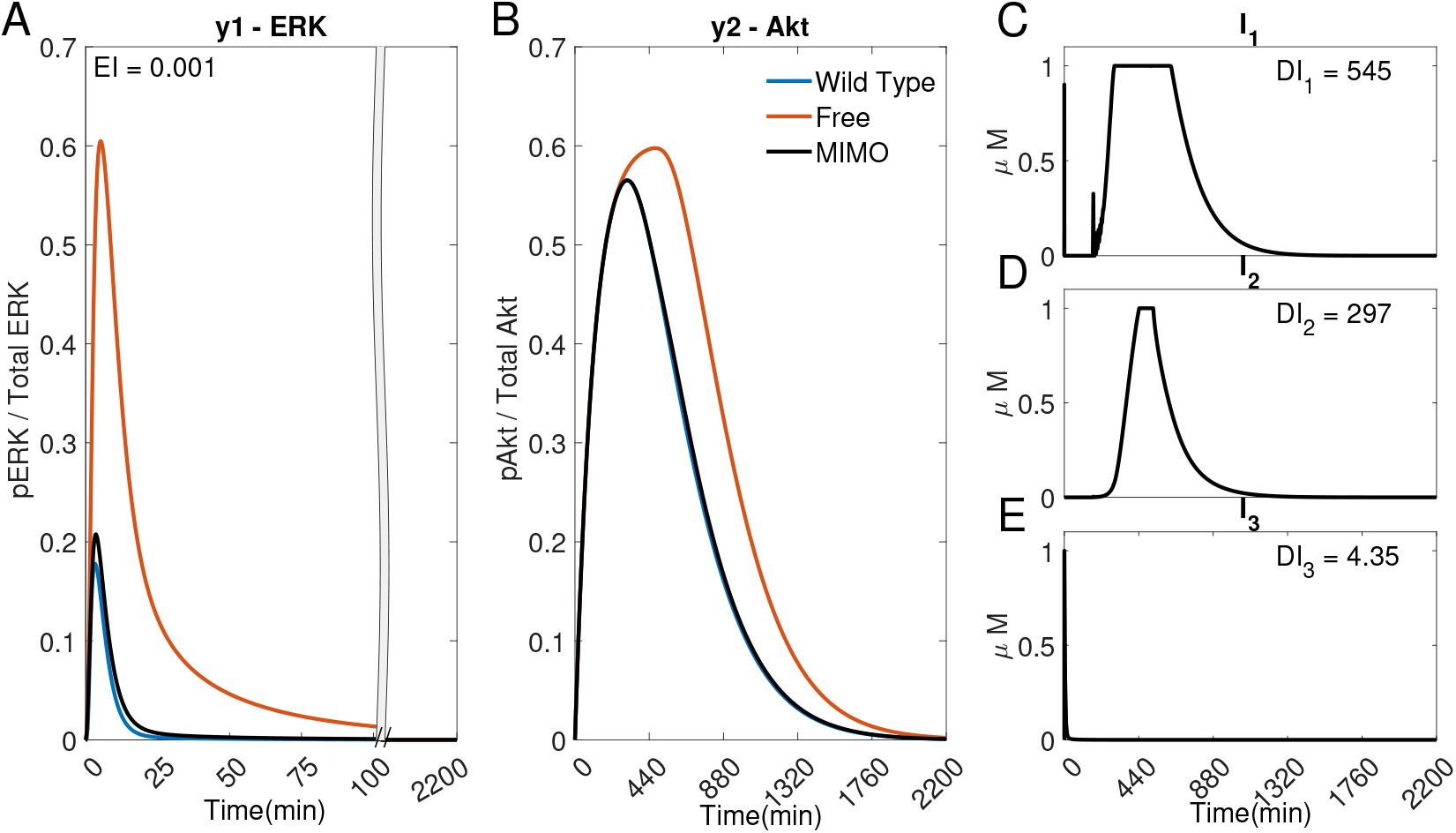
A MIMO adaptive MPC simulation using *I*_1_, *I*_2_ and *I*_3_ to control the concentrations of y1 (ERK) and y2 (Akt). (A) The response of ERK. (B) The response of Akt. (C-E) The inputs used in the simulation. Parameters: *T_s_* = 0.02*min, N* =10, *α* = 0, *β* = 0, *γ* = [1, 10^5^, 10^9^], *θ* = 0 and *η* =1.

To emulate capabilities of current microfluidics devices, the time step will be kept at *T_s_* = 1*min*. Therefore, in what follows, we investigate only Multi-Input Single-Output (MISO) simulations where y2 (Akt) is controlled only with *I*_1_ and *I*_2_. In this way, we remove the effect of the crosstalk (through the inherent negative feedback loops shown in Figure 1), which would restrict the MPC’s use of *I*_1_ and *I*_2_ caused by the high error in y1(ERK) due to the faster timescale of this output at the controller’s current time step (*T_s_* = 1*min*).

### 3.3 Combination Therapies Using *I*_1_ and *I*_2_

If cells are exposed to drugs for an extended period of time, side effects and resistance might become an issue [25]. The controller could be used to find potential drug combinations that can achieve a low Error Index (*EI*) whilst reducing the dose of the inputs (*DI_i_*). The weight associated with using each input, *γ*, within the cost function can be varied for this aim, as shown in Figure 7. The Bliss Independence (BI) formula [32,33] has been used here as a normalised Dose Index to summarise the combined effect of multiple drugs (see S5).

**Figure 7:**
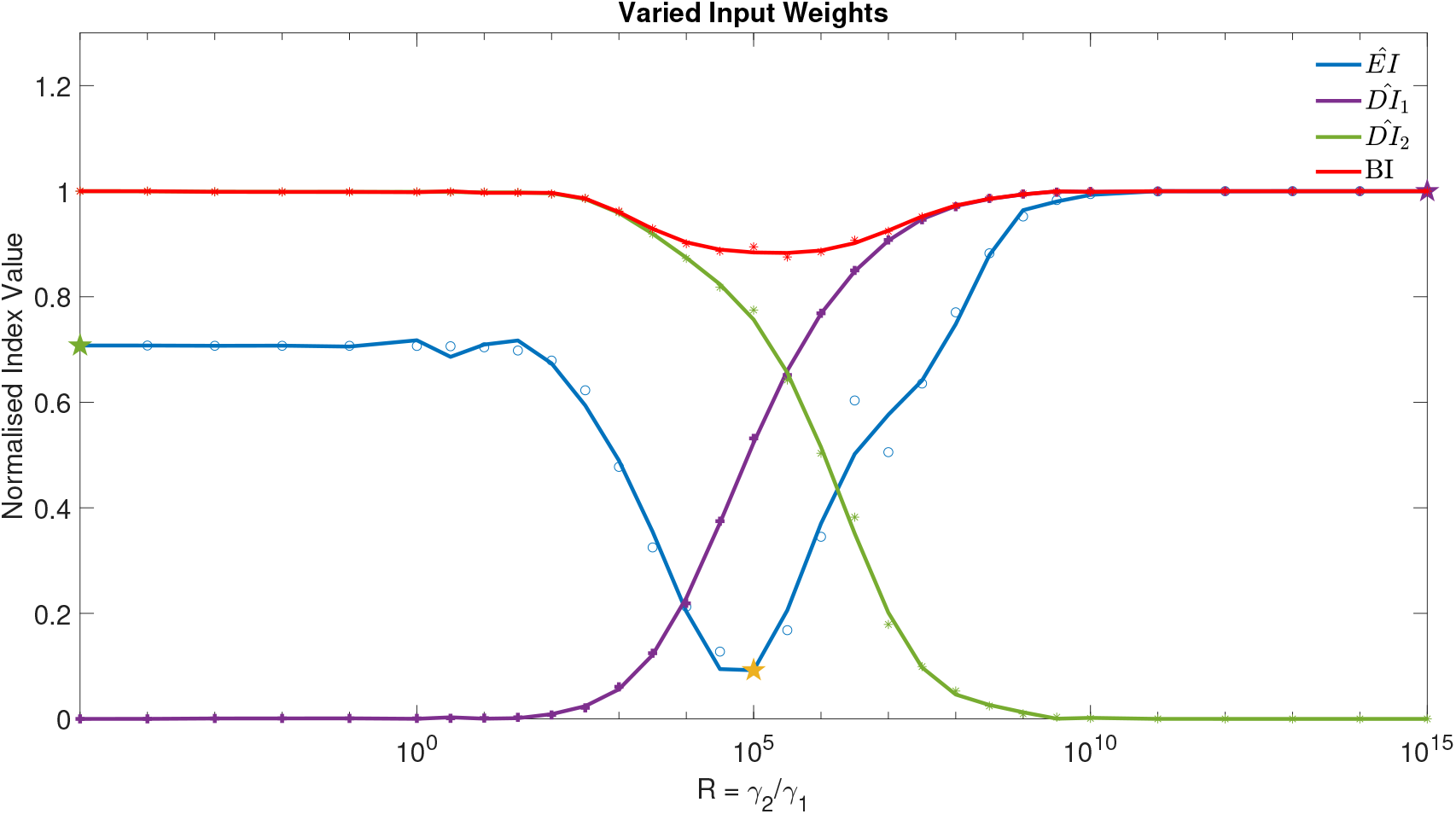
The normalised Error Index of y2 (Akt) ~, Dose Indexes of *I*_1_ ~ and *I*_2_ ~ and Bliss Independence ~, from 31 MISO adaptive MPC simulations using varied ratio of input weights, *R* = *γ*_2_/*γ*_1_, for example, when *R* = 100, *γ* = [1, 100, –]. Parameters: *T_s_* = 1*min*, *N* =10, *α* = 0, *β* = 0, *θ* = 0 and *η* =1. The three star markers show the ratio used in the plots of Figure 8

Figure 7 focuses on the control of y2 (Akt) using *I*_1_ and *I*_2_ as control inputs (MISO control). The weights in the cost function associated with each input can be varied as a ratio of 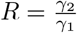, ranging from low *R* values (where a high weight is associated with *I*_1_, (*γ*_1_), thus producing a SISO-like simulation only using *I*_2_), all the way through to a high *R* value (where *γ*_2_ is relatively large and the controller will only use *I*_1_). Figure 7 compares the normalised Error Index, 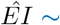, and the Bliss Independence *BI* ~ to the weight ratio (*R*). It shows that there is a range of *R* which can significantly reduce both *EI* ~ and *BI* ~. Therefore, the control performance of the MISO controller is better than any SISO simulation while keeping drug concentrations low. For the purpose of designing combination therapies, here the optimal input is associated to the minimum value of the *EI* ∼.

It can be seen from Figure 7 that the minimum occurs when *R* = 10^5^, corresponding to *γ*_1_ = 1 and *γ*_2_ = 10^5^. Figure 8 compares the responses obtained using a very low or high *R* value to the MISO simulation at the optimum of *EI*. This optimum achieves a significantly lower Error Index (*EI* → 0.25 ~ < 1.95 ~ < 2.75 ~), and a lower Dose Index (*DI*_1_ → 568 ~< 1068~ and *DI*_2_ → 423 ~ < 546~).

**Figure 8:**
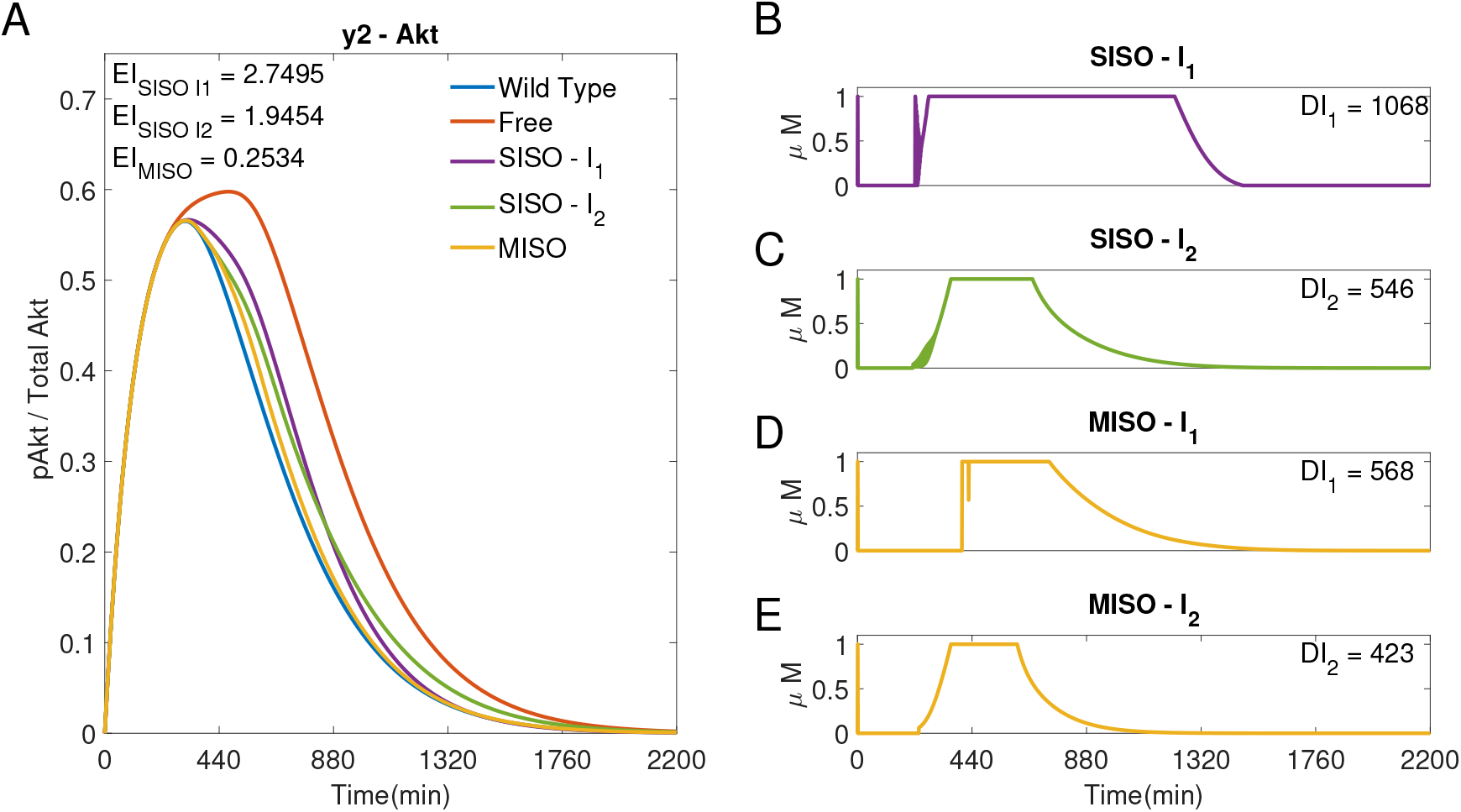
Three adaptive MPC simulations using *I*_1_ and *I*_2_ to control y2(Akt). (A) The response of y2 (Akt) to three different input weightings, SISO - *I*_1_ ~ (*γ* = [1, 10^15^, –]), SISO - *I*_2_ ~ (*γ* = [1, 10^−5^, –]) and MISO ~ (*γ* = [1, 10^5^, –]). *γ* was selected from the minimum point of the *EI* and the limits of *R* in Figure 7. (B, C) Inputs *I*_1_ and *I*_2_ used in the two SISO simulations. (D, E) Inputs *I*_1_ and *I*_2_, respectively, used in the MISO simulation. Parameters: *T_s_* = 1*min*, *N* =10, *α* = 0, *β* = 0, *θ* = 0 and *η* =1

### 3.4 Drug Holidays

If using an adaptive MPC, the user can set specific time intervals in which the controller does not give specific drugs (for example, to avoid toxicity induced by long exposure). These Drug Holidays can be achieved by the controller by varying the weights associated with each input, online, during a single simulation.

As an example, Figure 9 shows that the controller can retain a low Error Index whilst swapping inputs after 600 minutes (*EI* → 1.989 ≈ 1.950 < 2.75, the SISO *EI* of Figure 4). Therefore it is shown that a programmed change of cost function weights during the simulation can decide which input to stop using.

**Figure 9:**
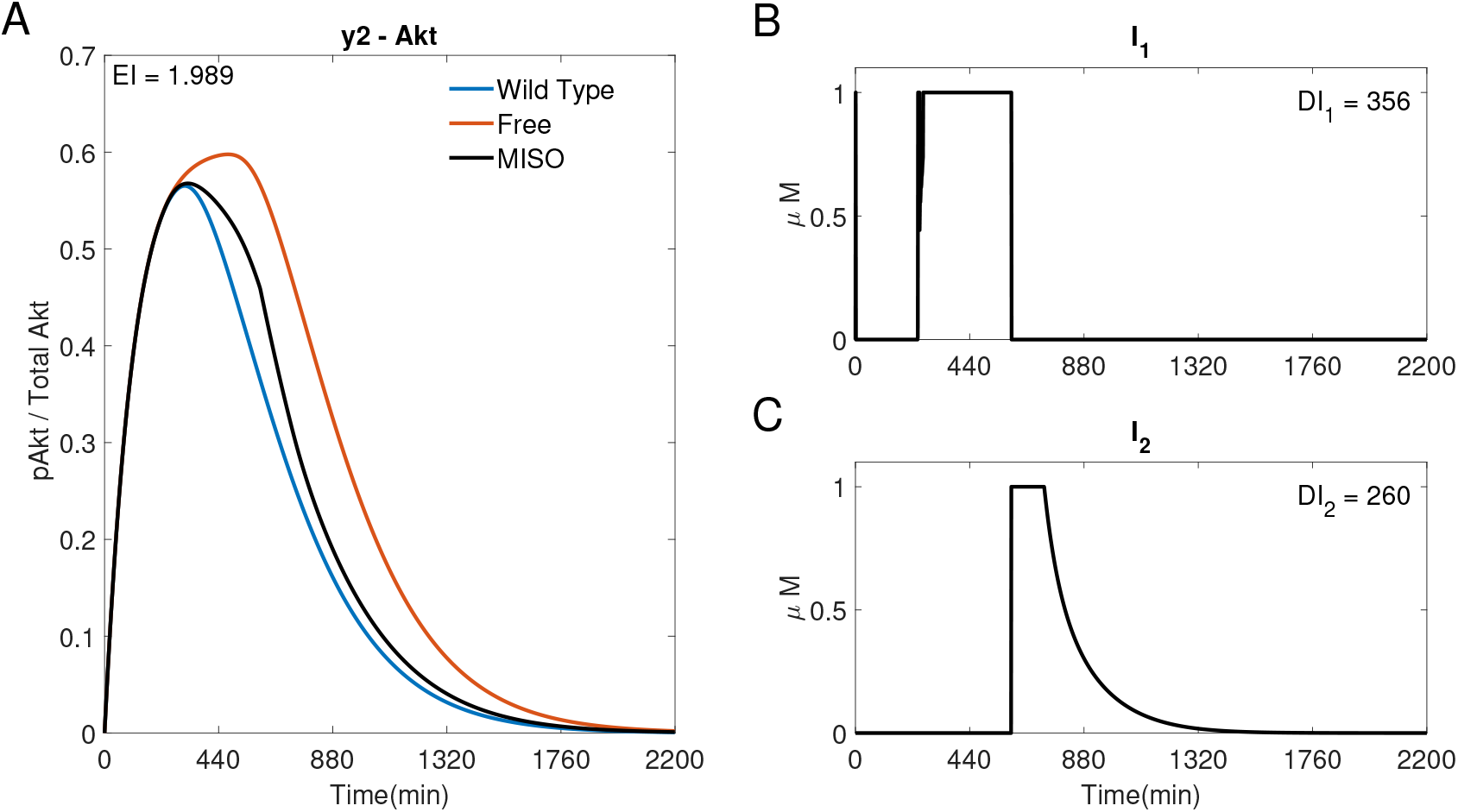
A MISO adaptive MPC simulation swapping *I*_1_ and *I*_2_ to control y2 (Akt). (A) The response of Akt. (B, C) The inputs used in the simulation. Parameters: *T_s_* = 1*min*, *N* =10, *α* = 0, *β* = 0, *γ* = [1, ∞, –] when 0 ≤ t ≤ 600min and *γ* = [∞, 10^5^, –] when 600 < *t* ≤ 2200min, *θ* = 0 and *η* =1.

Alternatively, the controller can be set to only choose one input at each time step. The inputs shown in Figure 10 have an ON or OFF state, 1*μM* or 0*μM* (discrete inputs). The step size, *T_s_*, used makes a significant difference to the response. When *T_s_* = 1*min* the drugs can switch ON or OFF every minute, leading to rapidly fluctuating inputs (shown in Section S6). Figure 10 shows the discrete simulation with a larger time step (*T_s_* = 30*min*). It can be seen that there is a better performance when compared to the SISO simulations (Figure 4) as *EI* → 1.6878 < 1.95 < 2.75, whilst still using less of each input (*DI*_1_ → 630 < 1068, *DI*_2_ → 450 < 546). However, when compared to the optimal MISO response (Figure 8), these added constraints result in a higher Error Index (*EI* → 1.6878 > 0.2534) and a higher dose (*DI*_1_ → 630 > 568, *DI*_2_ → 450 > 423).

**Figure 10:**
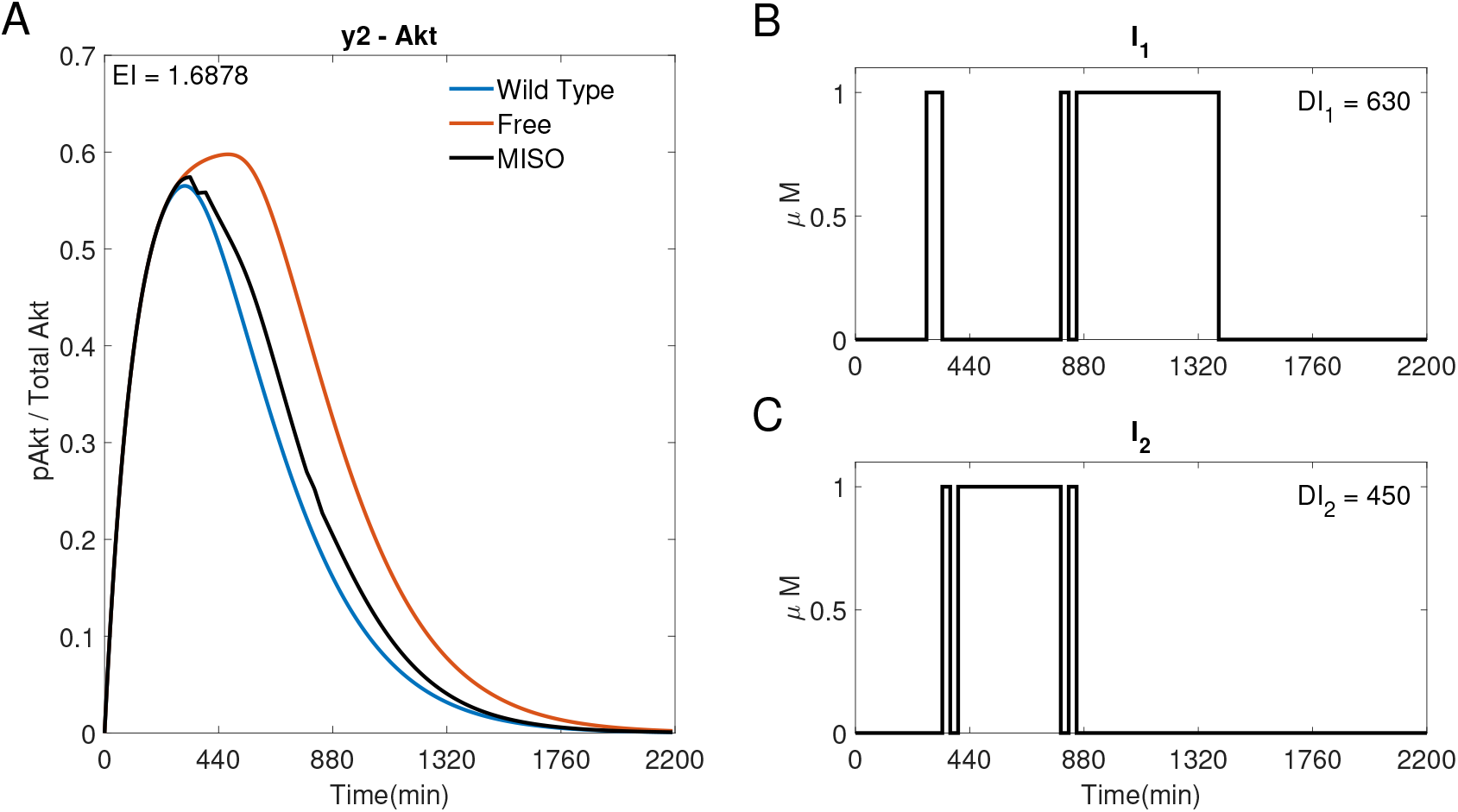
A discrete MISO adaptive MPC simulation using *I*_1_ and *I*_2_ to control y2 (Akt). (A) The response of Akt to the inputs in (B) and (C). Parameters: *T_s_* = 30*min*, *N* =10, *α* = 0, *β* = 0, *γ* = [1, 10^5^, –], *θ* = 0 and *η* =1.

## 4 Discussion

Computational methods have been extensively used in the search for effective cancer treatments, with approaches including optimal control to regulate dynamics of different cell populations [34–43], and a feedback action to account for changes in the cancer system [44–51]. This is, to the best of our knowledge, the first attempt of using feedback control to regulate intracellular dynamics in cancer cells.

We showed that an adaptive MPC program can be used to inform treatments for NSCLC cells, steering the dynamics of several key signalling pathways, whilst offering a tunable cost function that allows to adjust the characteristics of an optimal input. Indeed, the controller can be tuned to choose different drug profiles that will achieve a similar control performance whilst reducing exposure to one or more drugs.

Other control strategies, like PID controllers, cannot be directly tuned to effect the desired input and only act on the observed output. The use of a linear model within the MPC algorithm makes the control algorithm running time short enough for it to be used, in the future, in external feedback control experiments. The implementation of those would require some practical aspects to be considered, which we did not account for. Firstly, there might be delays in cell responses to drugs/actuation, which the model used by the controller should account for. Also, the sampling/actuation time might need to be fast enough, if aiming at controlling genes with fast dynamics. This issue might be overcome using experimental optogenetics-based platforms instead of microfluidics-based ones, as they can reduce delays in the actuation. Finally, this method assumes the knowledge of a model. In the future we hope to work on an controller that adapts the model based on online system measurements.

We foresee a growing interest in applying cybergenetics approaches, and in particular feedback controllers, to steer mammalian cells dynamics. If we realise our ambition to implement the experiments proposed here on living cells and, longer term, on patient-derived organoids, feedback control might be a valuable tool for the design or personalised optimal treatments for a range of conditions.

## 5 Conclusions

It has been demonstrated that adaptive MPC can be used to inform treatments for NSCLC cells, guiding the behaviour of two key signalling pathways and offering a tunable cost function to modify *ad hoc* the rapidity of inputs’ changes, and the ways inputs as chose. The weights can also be changed during its operation to choose different drug profiles that will achieve a similar control performance whilst giving the cell a break from individual drugs. In the future, we hope to test the controller in living cells using microfluidics/microscopy platforms.

## 6 Author Information

### 6.1 Author Contributions

B.S. designed and carried out the simulations in Matlab2021b and wrote the manuscript with inputs from all the authors. I.d.C. supported the project. L.R. and L.M. conceived the project, contributed to manuscript writing and supervised the entire work.

### 6.2 Declaration of Interests

Nothing to declare.

## 7 Acknowledgements

B.S. is supported by an EPSRC DTP Scholarship. L.R. is funded by a Research Fellowship from the Royal Academy of Engineering (RF1516/15/11). L.M. is supported by the Engineering and Physical Sciences Research Council (EPSRC) grant EP/S01876X/1, and by the European Union’s Horizon 2020 under Grant Agreement No. 766840.

## S1 NSCLC Model

The model of the NSCLC is [27]

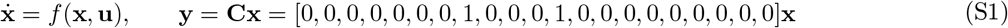

where the vector field *f*(.,.) is detailed in (S4)–(S23), **x** is the state vector containing 21 molecule concentrations (Table S1), **u** is the input vector **u** = [*I*_1_, *I*_2_, *I*_3_]^*T*^ (orange in (S4)–(S23), each input acts on both the active and inactive target molecule and therefore appears twice in (S4)–(S23)) and **y** is the vector of outputs *y*_1_ = *pERK* and *y*_2_ = *pAkt* (blue in (S4)–(S23)).

**Table S1.**

| $\mathbf{x}_i$ | Shorthand | Molecule |
| --- | --- | --- |
| $x_1$ | pEGFR | Active epidermal growth factor receptor |
| $x_2$ | DSOS | Deactive SOS |
| $x_3$ | SOS | Son Of Sevenless, a Guanine nucleotide exchange factor |
| $x_4$ | Raf | Raf kinase |
| $x_5$ | pRas | Active Ras, a small GTPase |
| $x_6$ | pMEK | Active Methyl ethyl ketone |
| $x_7$ | ERK | Extracellular-signal-regulated kinase |
| $x_8$ | pERK | Active ERK |
| $x_9$ | pIGF1R | Active Insulin-like growth factor receptor |
| $x_{10}$ | PI3K | Phosphoinositide 3-kinase |
| $x_{11}$ | pPI3K | Active PI3K |
| $x_{12}$ | pAkt | Active Akt |
| $x_{13}$ | Akt | Set of three serine/threonine-specific protein kinases |
| $x_{14}$ | PP2A | Protein phosphatase 2, Kinase inhibitor |
| $x_{15}$ | Ras | A small GTPase |
| $x_{16}$ | pRaf | Active Raf |
| $x_{17}$ | MEK | Methyl Ethyl Ketone |
| $x_{18}$ | RasGAP | GTP hydrolyser of Ras |
| $x_{19}$ | ppRaf | Raf which has been phosphorylated twice |
| $x_{20}$ | P90 | Ribosomal S6 kinase |
| $x_{21}$ | pP90 | Active P90 |

### S1.1 Difference Between NSCLC and Wild Type Cells

Mutations present in NSCLC cells can lead to an overexpression of *EGFR* and *IGF*1*R*; this is represented by using different initial conditions for *pEGFR* and *pIGF*1*R* [27] and leads to the response shown in Figure 2.

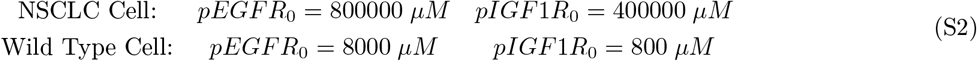

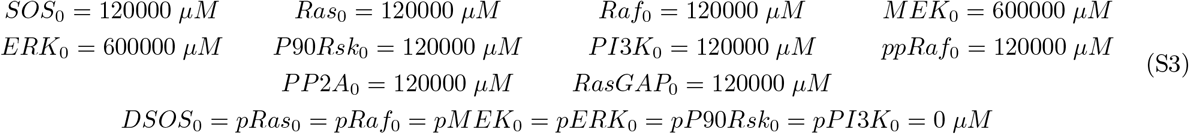

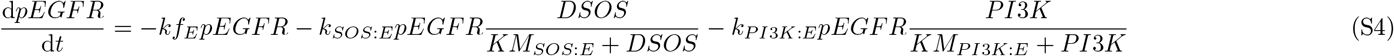

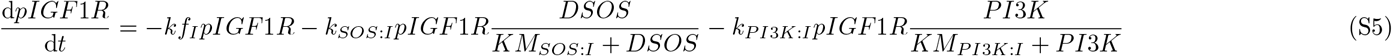

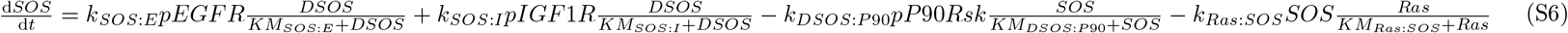

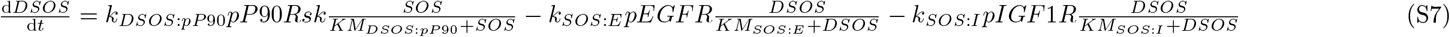

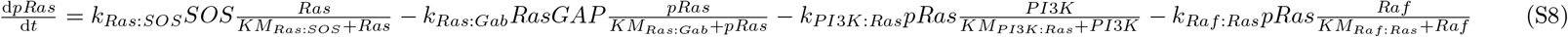

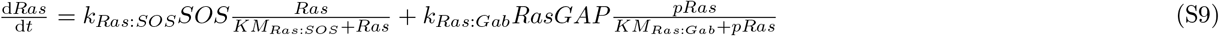

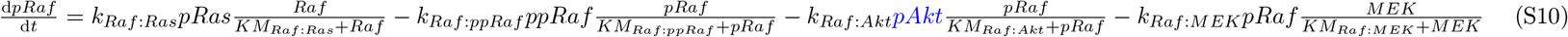

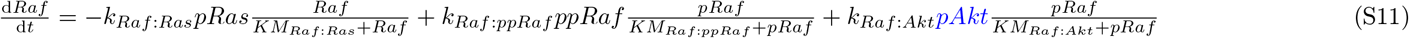

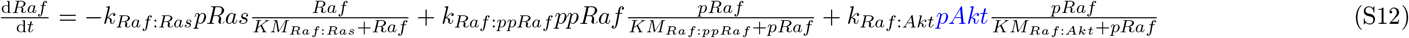

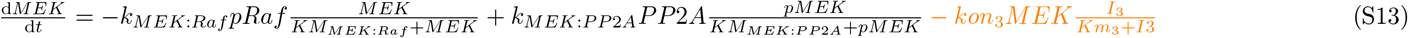

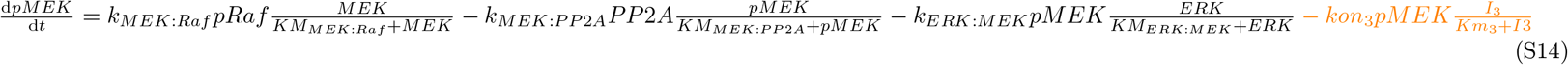

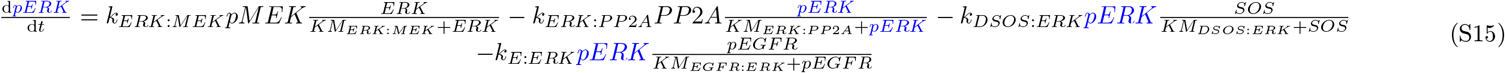

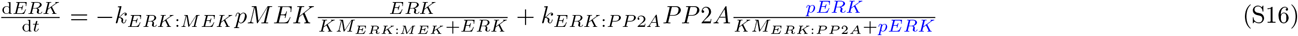

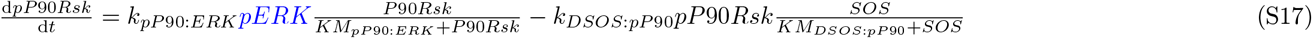

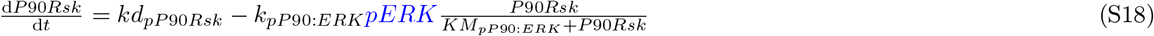

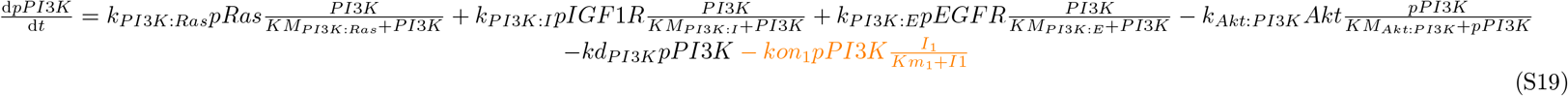

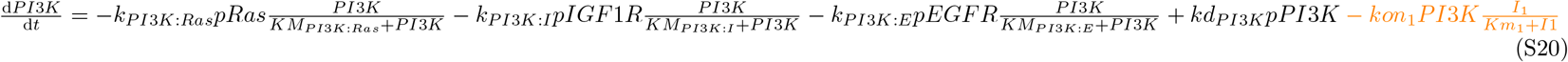

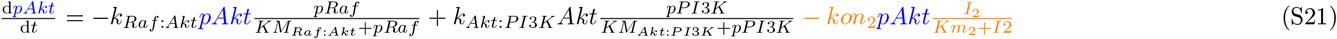

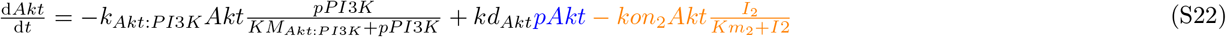

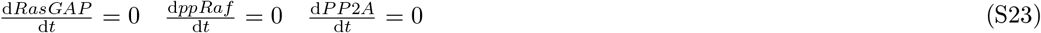

### S1.2 Conservation Equations

Each molecule can either be active, *p*(.), or inactive, but there is a constant total concentration of each molecule within the model. This total, (.)_*T*_, is defined by the following conservation equations:

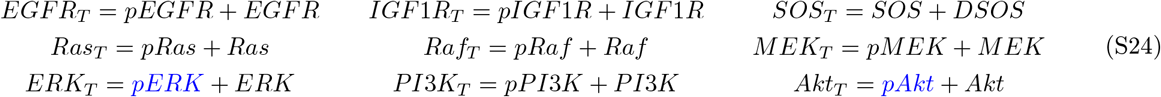

### S1.3 Parameters in the NSCLC Model

**Table S2:**
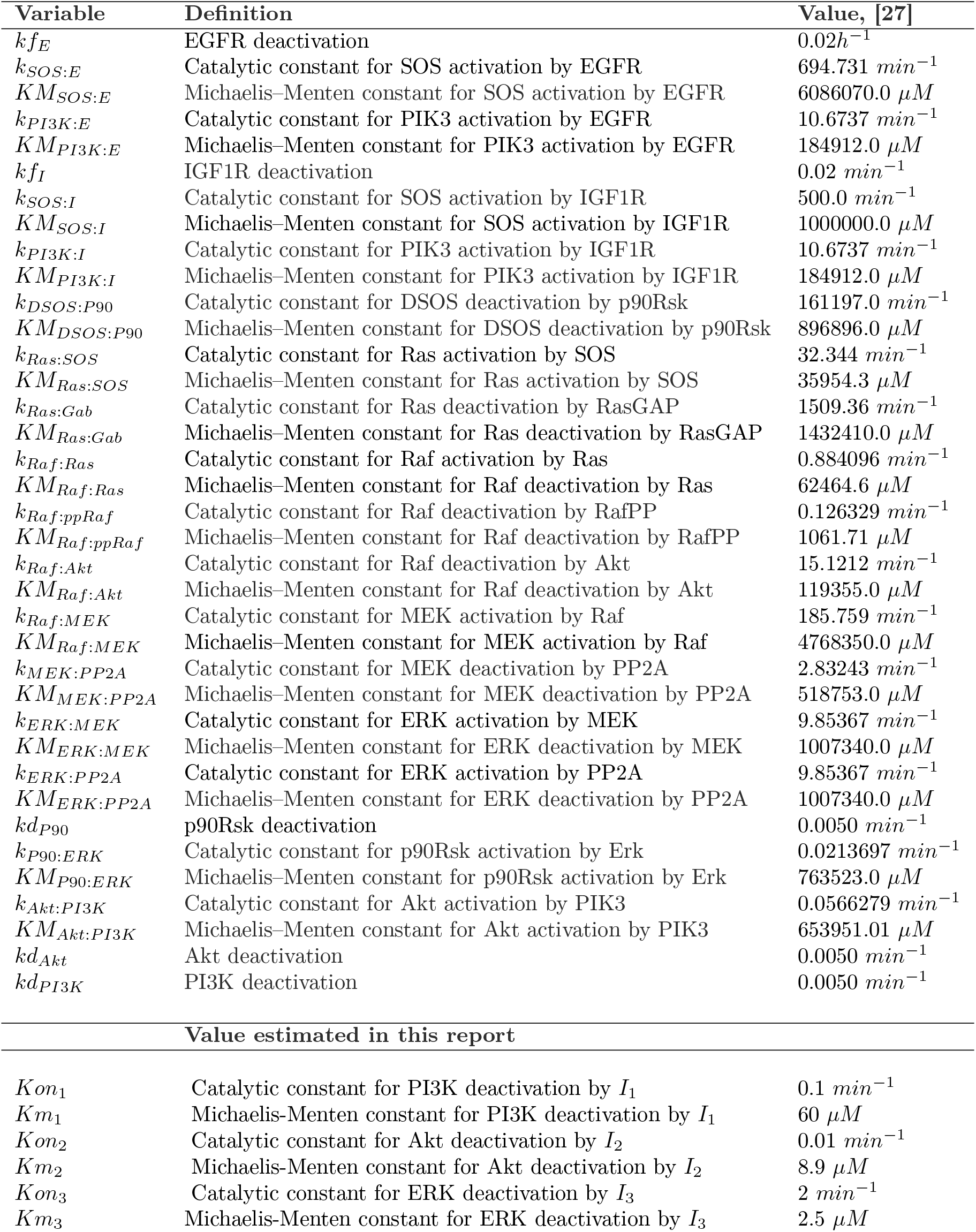
Parameters used in the NSCLC model including the six input parameters discussed in Section S2.

## S2 Parameter Choice For Input Interactions

The equation governing the inputs reaction with the target molecule is given by

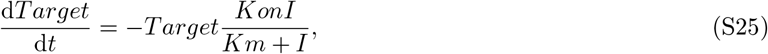

where the Michealis-Menten parameter values associated with each input are reported in Table S3. *Km* is equivalent to the *IC*_50_ value of the inhibitor being used on each target. To make the response of the controller inputs less ‘switch like’, inhibitors with relatively large *IC*_50_ values have been chosen. *K on* has then been chosen by fitting the model simulations to western blots for *I*_1_-3MA [52], *I*_2_-Oridonin [53] and *I*_3_-Pimasetib [54].

**Table S3:** Input properties used in the simulations

| Drug | $Km(\mu M)$ | $Kon(min^{-1})$ | Target Molecule |
| --- | --- | --- | --- |
| $I_1$ - 3MA | 60 [55] | 0.1 | PI3K |
| $I_2$ - Oridonin | 8.9 [56] | 0.01 | Akt |
| $I_3$ - Pimasetib | 2 [57] | 2.5 | MEK |

## S3 Step Input Response

Step input simulations for each input, as described in Section 3.1.1, can be seen in Figure S1. In all cases, the control performance is worst as compared to that obtained with SISO control (Figure 4), as the *EI* is either comparable (S1A) or higher (S1C and E) than those obtained applying feedback control; also, in all step input simulations, the Dose Index is higher.

**Figure S1:**
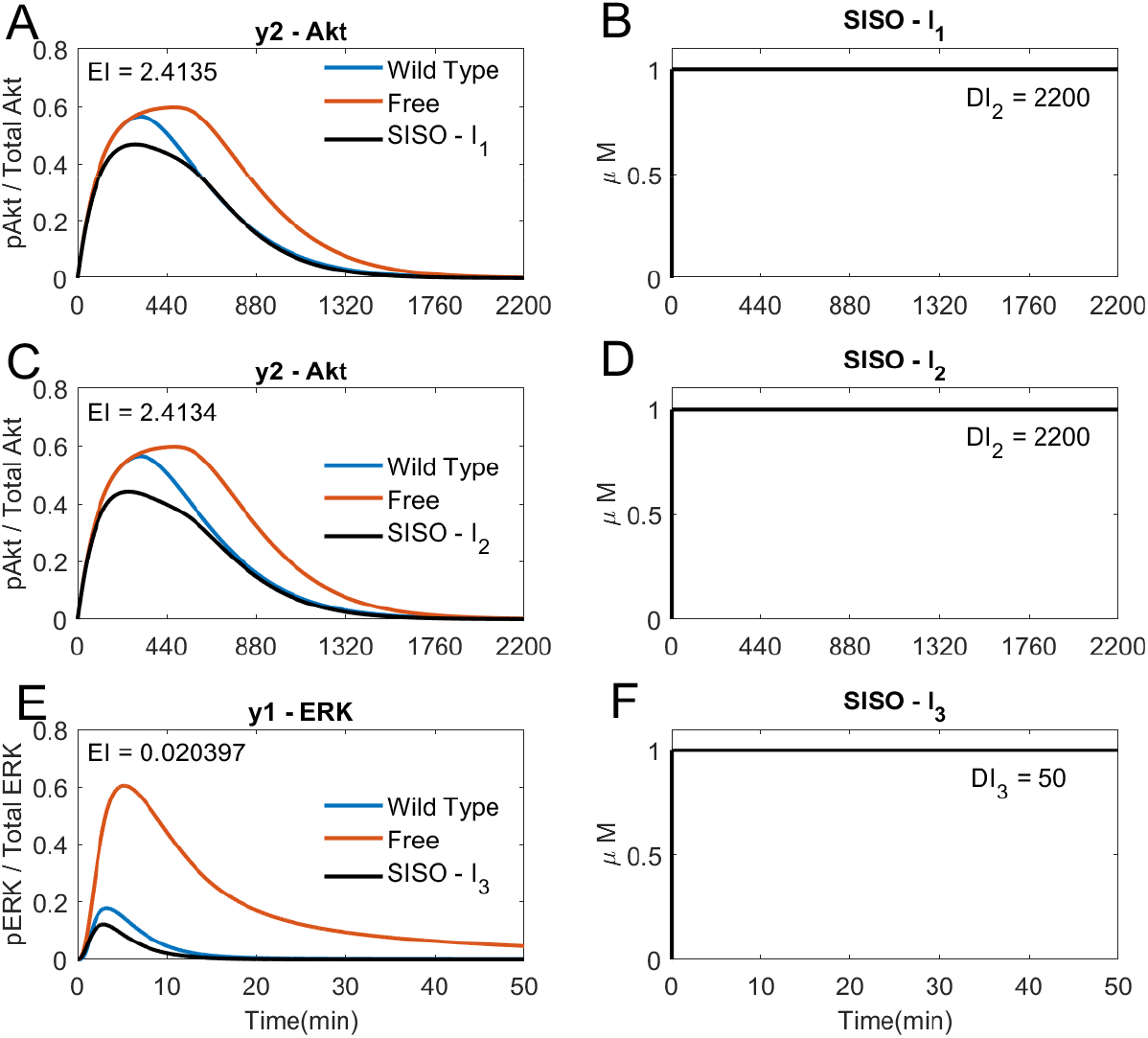
Step Input simulations. (A) The response of y2 (Akt) to the step input of *I*_1_ in (B). (C) The response of y2 (Akt) to the step input of *I*_2_ in (D). (E) The response of y1 (ERK) to the step input of *I*_3_ in (F).

## S4 Model Predictive Control (MPC)

MPC uses a model of the plant (system to be controlled) in the feedback loop to estimate the effect of the inputs the controller will choose at each time step. The inputs are chosen to minimise a user-defined cost function that typically includes terms penalising the magnitude of the inputs, **u**(*t*), and the magnitude of errors, **e**(*t*), between the response of the system and reference signals [29], as shown in Figure S2. The feedback loop then measures the outputs of the actual plant, **y**(*t*), to estimate the actual states, **x**(*t*), and reiterates the MPC scheme to choose each subsequent input, **u**(*t*).

Some modifications to the cost function are discussed in Sections S4.2 and S4.3. MPC controllers are not limited to linear systems, however, non-linear systems can result in a larger computational effort and require more complex optimisation solvers, as discussed in Section S7.

**Figure S2:**
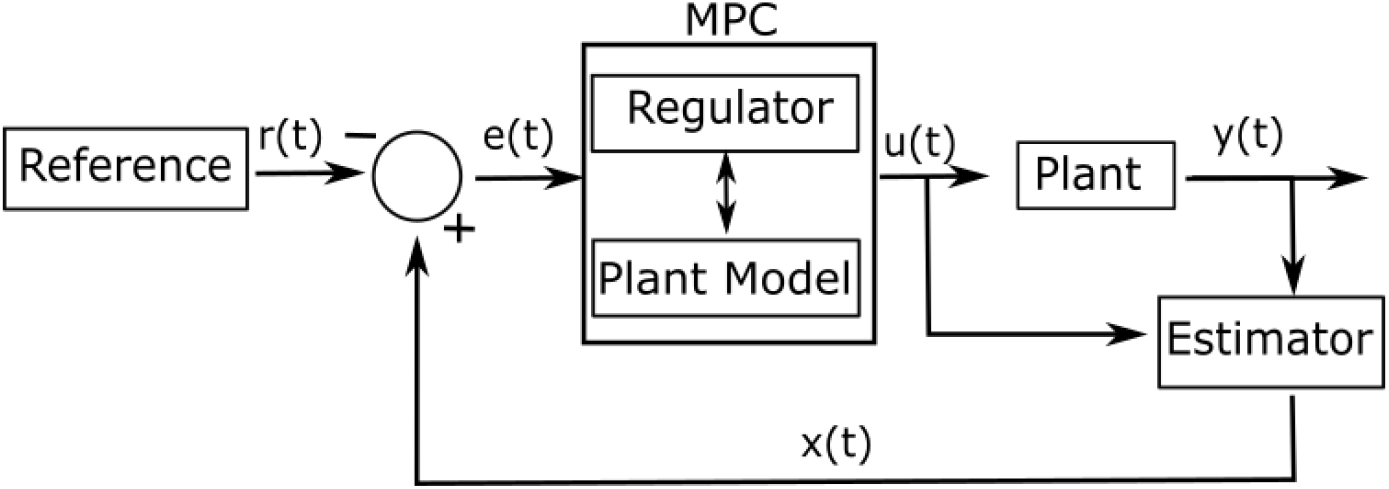
MPC architecture.

For the present numerical study, the plant is the nonlinear NSCLC model presented in Equations (S4)–(S23) and all the system states, **x**(*t*), are measured directly. In practice, only a few outputs/states can be measured and an estimator is required to estimate the remaining states from input-output data as shown in Figure S2.

The control reference signal, **r**(*t*), is chosen as the response of a wild type cell (i.e. without cancer). **e**(*t*) is the error signal between the reference, **r**(*t*), and internal states of the plant, **x**(*t*). **e**(*t*) is fed into the regulation block of the control scheme; this is where the optimisation problem is solved.

### S4.1 Cost Function Derivation

The internal state errors **e**(*t*) are calculated and fed into the MPC block at each time step. The MPC controller uses the model of the plant system to predict the future state error of the system for possible combinations of inputs, within the problem constraints, over the prediction horizon, *N*. The controller then optimally chooses the input profile that results in the minimum of a predetermined cost function, *J*(**U**), in the regulator. The inputs of the first time step of this optimal sequence are then applied to the plant system. At the next time step, the error in the states is estimated and this process repeats.

Usually, the optimisation problem contains the cost function to be minimised, and the state and input constraints.

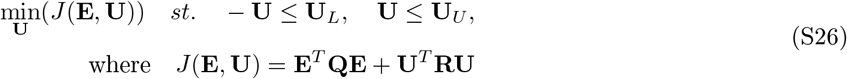

The model that the regulator sees is a discrete approximations of the NSCLC model for 1 ≤ *k* ≤ *N* steps. **E** = [**e**(0), **e**(1), …, **e**(*N*)]^*T*^ describes the current and future predicted state errors. **U** = [**u**(1), **u**(2), …, **u**(*N*)]^*T*^ describes the future inputs. The weight of the cost function related to each term can vary what is considered the optimal input. **Q** weights the error of the states and **R** weights the use of the inputs [29]. For example, if we want to minimise the use of a drug then the weight of the cost function associated with the inputs, **R**, should be relatively large compared to the weight of the state error, **Q**. Constraints on the input bounds are also included here (**U**_*L*_ ≤ **U** ≤ **U**_*U*_, in all simulations **U**_*L*_ = 0*μM* and **U**_*U*_ = 1*μM*).

#### S4.1.1 Linear MPC

If a linear approximation of the model can be produced and represented by a state space, the future behaviour of the model can be calculated offline and the optimisation problem is convex (for affine constraints). The cost function can be minimised using a quadratic solver, which is relatively computationally light (MATLAB R2021b’s ‘quadprog’ solver was used here). Here, a state space was used to represent the plant’s model:

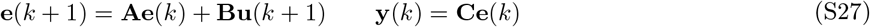

As long as the current state is known, each future state can be estimated for a given input profile, i.e.

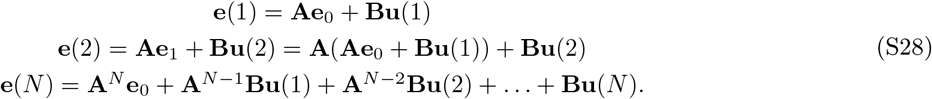

Defining a new notation containing all of the states in the prediction horizon we get:

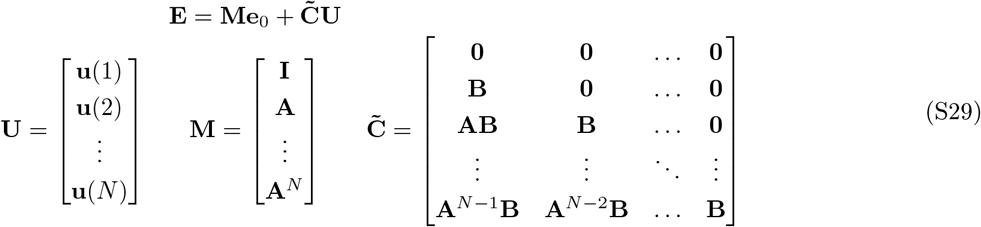

Therefore, the cost function can be rearranged as

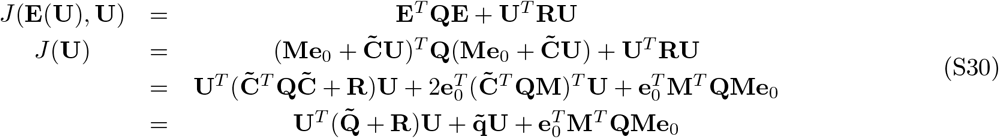

The final term in the cost function can be removed as it is constant with respect to the inputs and therefore will not affect the position of the minimum point; thus,

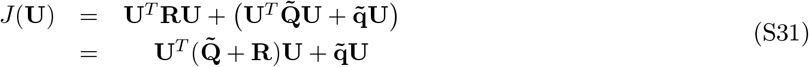

Only the optimal inputs for the first time step are then applied to the plant, **u**(1), the whole optimisation process is repeated at the next time step.

#### S4.1.2 Weighting of the Traditional Cost Function

The cost function as shown in (S30), can be weighted as a balance of using the inputs, *γ*; the error in all of the estimated states, *α* and the error in the outputs, *β*.

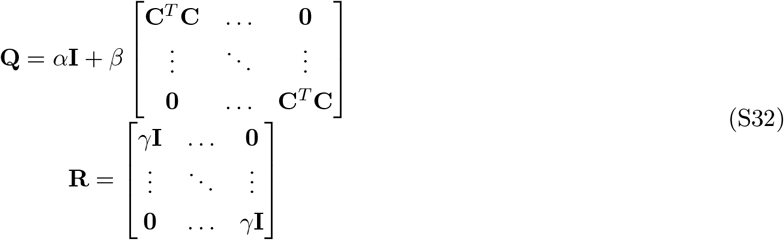

where **I** is an identity matrix. The cost function can be tuned by *α*, *β* and *γ*.

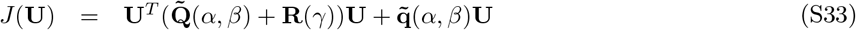

### S4.2 Differential Terms in the Cost Function

The controller can favour to rapidly change input concentrations, which is not ideal for *in vitro* experiments, where there might be delays in the actuation, and frequent media change might cause stress to cells. Therefore, a term related to the gradient of the inputs was added to the cost function to reduce fast variations of the inputs. The linear approximation of the model used within the MPC simulations is discrete and therefore the derivative is approximated by the scaled difference between inputs at adjacent time steps.

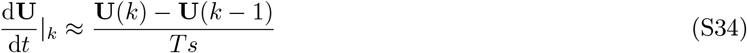

Using the squared sum of the derivative of the inputs, the gradient of the steps between the last actual input and the last step in the prediction horizon can be added to the cost function. Any constant scaling can be dropped as this would just change the weighting added to the term.

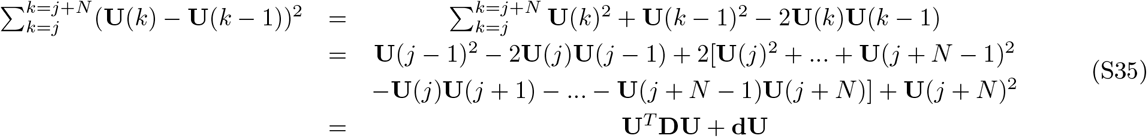

The gradient can be added to the cost function, *J*(**U**), (Equation S33,) through **D** and **d**, pre-multiplying by *θ*.

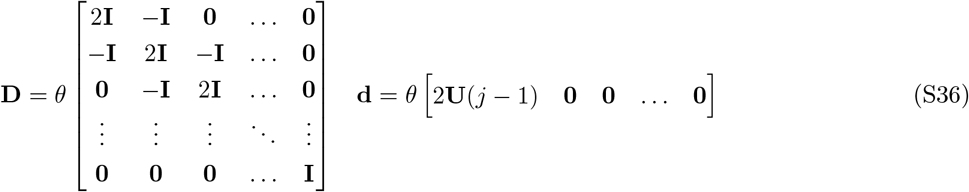

where **I** is a square identity matrix (size equal to the number of inputs).

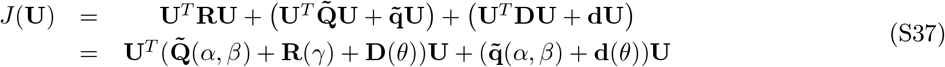

Figure 3 demonstrates the effect of the differential cost.

An alternative approach would have been to add the gradient as a constraint into the optimisation problem preventing the inputs from varying faster than a limiting value. However, this would have merely limited the maximum rate of inputs’ variation.

### S4.3 Integral Terms in the Cost Function

When controlling a signalling phosphorylation cascades, it is desirable to decrease both the peak and the duration of an error [30]. Both the duration and the peak are included in the integral of the outputs. Therefore the integral of the output errors should be added as a term in the cost function. The square of the integral errors has been approximated.

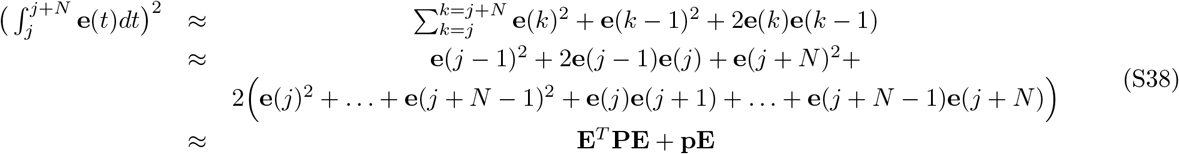

The first term is constant and is therefore dropped. **P** and **p** are weighted by *η* in the cost function.

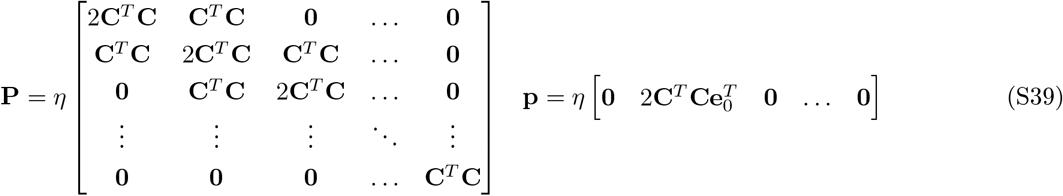

The integral cost is defined in terms of the future state errors **E** rather than the inputs **U**. The matrices defined in Equation (S29) can be used.

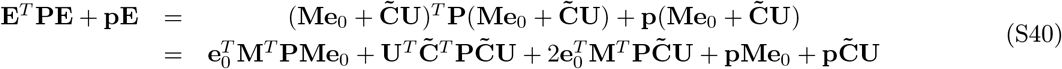

The constant terms with respect to **U** can be dropped and the quadratic and proportional terms reorganised.

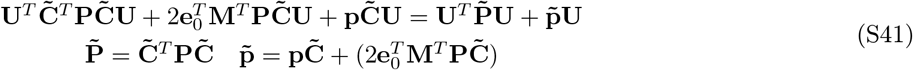

The cost function, *J*(**U**) (from Equation S37), can be formed including the integral of the state error.

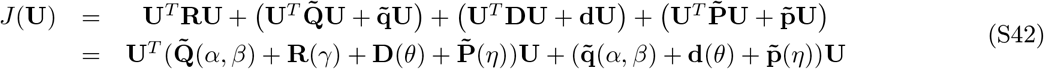

This is the cost function that has been used in all the simulations with different weights. Figure 3A) demonstrates the effect of the integral cost on reducing both the amplitude and the duration of the outputs.

### S4.4 Adaptive MPC

Adaptive MPC describes an MPC control scheme in which the model of the plant changes as the simulation progresses [58]. Here, it is assumed that the states of the actual plant can be perfectly measured at each step (as no estimator is used). Our non-linear model of the NSCLC, (S4) - (S23), can then be linearised about these current measurements of the states, **x**(*k*), at each time step. This linear model forms the state space in Equation S27 and the cost function is reformed at each iteration to better represent the local future dynamics of the plant. The adaptive linear MPC has a better performance than a single linear model and is less computationally expensive than using a full non-linear model [28,59,60].

## S5 Normalisation of Indexes And Bliss Independence

In order to easily compare simulations, it is useful to have an index which summarises the performance of the controller and the type of input it takes to achieve this performance, *EI* and *DI_i_*, respectively (normalised here such that multiple plots can be compared within Figure 7).

The Error Index, *EI*, is the sum of the squared errors of the outputs, calculated by integrating the square of all the output error signals by using a trapezium approximation of the discrete data:

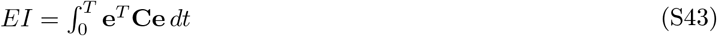

The largest *EI* (worst performance) found in any MPC simulation using the chosen cost function to control y2 (Akt) is the SISO simulation using *I*_1_ (*EI* = 2.75, Figure 4). This has been used to normalise *EI* to form 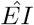.

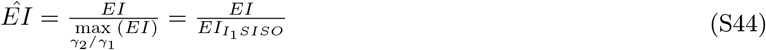

*DI_i_* is equivalent to the integration of input profile for each input, *I_i_*:

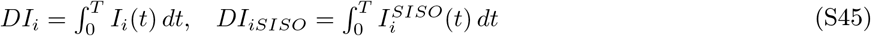

*DI_i_* is normalised by the *DI_i_* of *I_i_*’s SISO simulation (*I*_1_ and *I*_2_ acting on y2 (Akt) and *I*_3_ acting on y1 (ERK) in Figure 4, *DI*_1*SISO*_ = 1068, *DI*_2*SISO*_ = 546, *DI*_3*SISO*_ = 4), producing 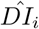.

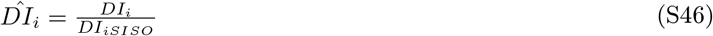

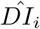 does not give a quantitative measure on the dose of the combined input profile. Within current literature, there are many methods of trying to summarises the joint effect and toxicity of combination therapies, where multiple drugs are given together at a determined time point [32]; however these do not look into dynamic dosages over a given time period. Therefore a combined effect of the drug profiles can be estimated by replacing these static drug dosages with the normalised Dose Index, 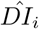.

An Isobole can be defined as 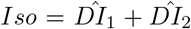 for these therapies. From this definition our combinations are all antagonistic. The Bliss Independence formula, *BI*, assumes that there is no correlation between the two agents.

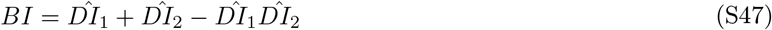

Our model is deterministic, with each input having a different target molecule, therefore within these *in silico* simulations there is no correlation between the inputs. Therefore the Bliss Independence formula can be used to gauge the combined effect of 2 drugs [33]. All three indexes are used in Figure 7 to compare multiple simulations.

## S6 Discrete Drug Holiday

A discrete MISO simulation, as described in Section 3.4, where *T_s_* = 1*min* can be seen in Figure S3. Figure S3 achieves an *EI* = 1.3533, significantly better than just a SISO simulation in Figure 4 (*EI* → 1.35 < 1.95 < 2.75) whilst using a lower Dose Index (*DI*_1_ → 645 < 1068 and *DI*_2_ → 438 < 546). However in this simulation both inputs rapidly fluctuate from 0*μM* to 1*μM*, therefore a longer time step can be used to reduce the fluctuations.

**Figure S3:**
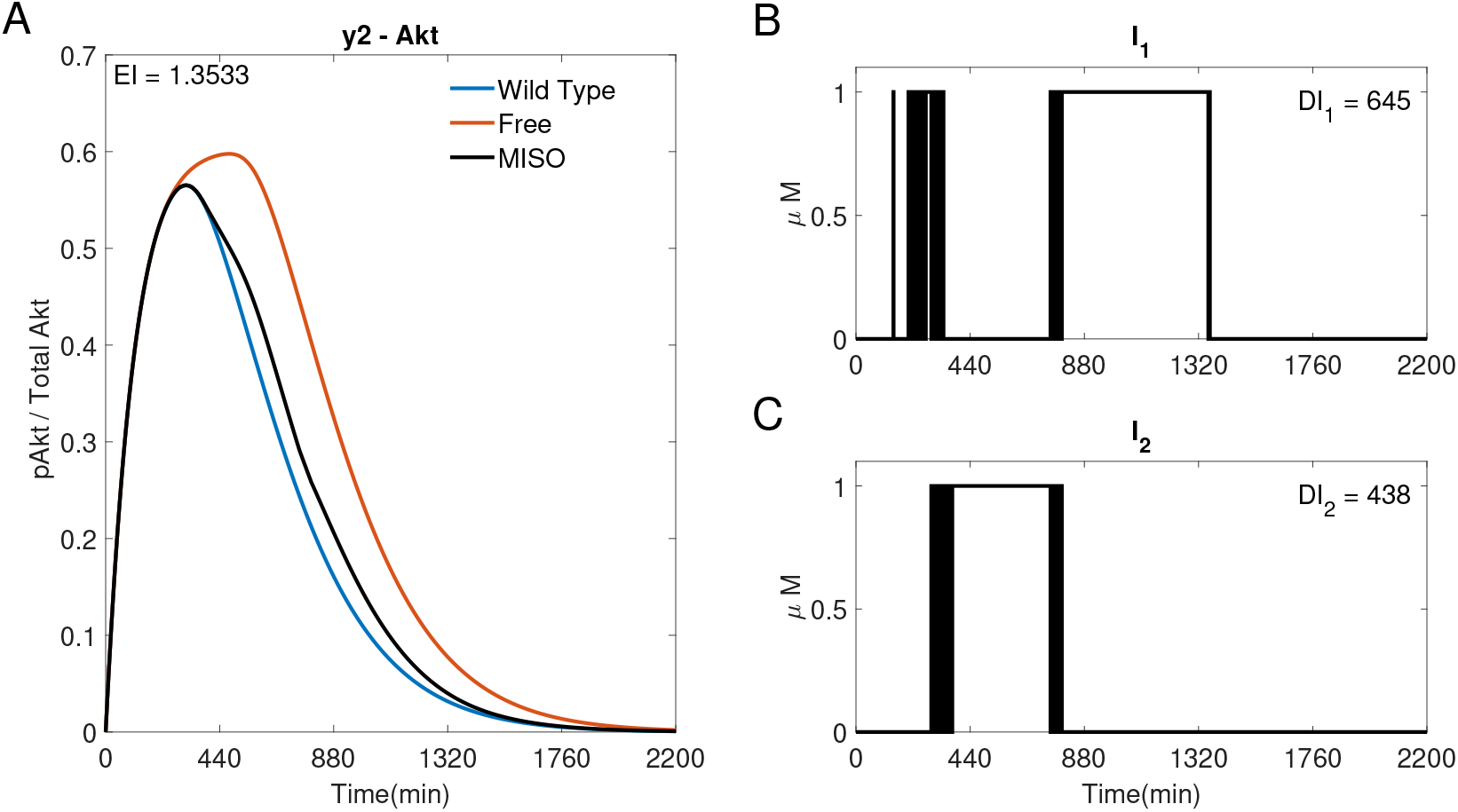
A discrete MISO adaptive MPC simulation using *I*_1_ and *I*_2_ to control the concentration of y2(Akt). A) The response of Akt. B) and C) The inputs used in the simulation. Parameters: *T_s_* = 1*min*, *N* =10, *α* = 0, *β* = 0, *γ* = [1, 10^5^, –], *θ* = 0 and *η* =1.

## S7 Linear vs Non-Linear MPC

All MPC simulations use an adaptive linear MPC controller, where the linear model is based off a linearisation of a non-linear model of the NSCLC system (S4)–(S23). Using non-linear MPC creates a non-convex optimisation problem requiring a more complex (and computationally heavy) solver. The non-linear simulations have used MATLAB’s ‘fmincon’, a gradient based non-linear solver. It is the fastest appropriate solver in MATLAB R2021b.

**Figure S4:**
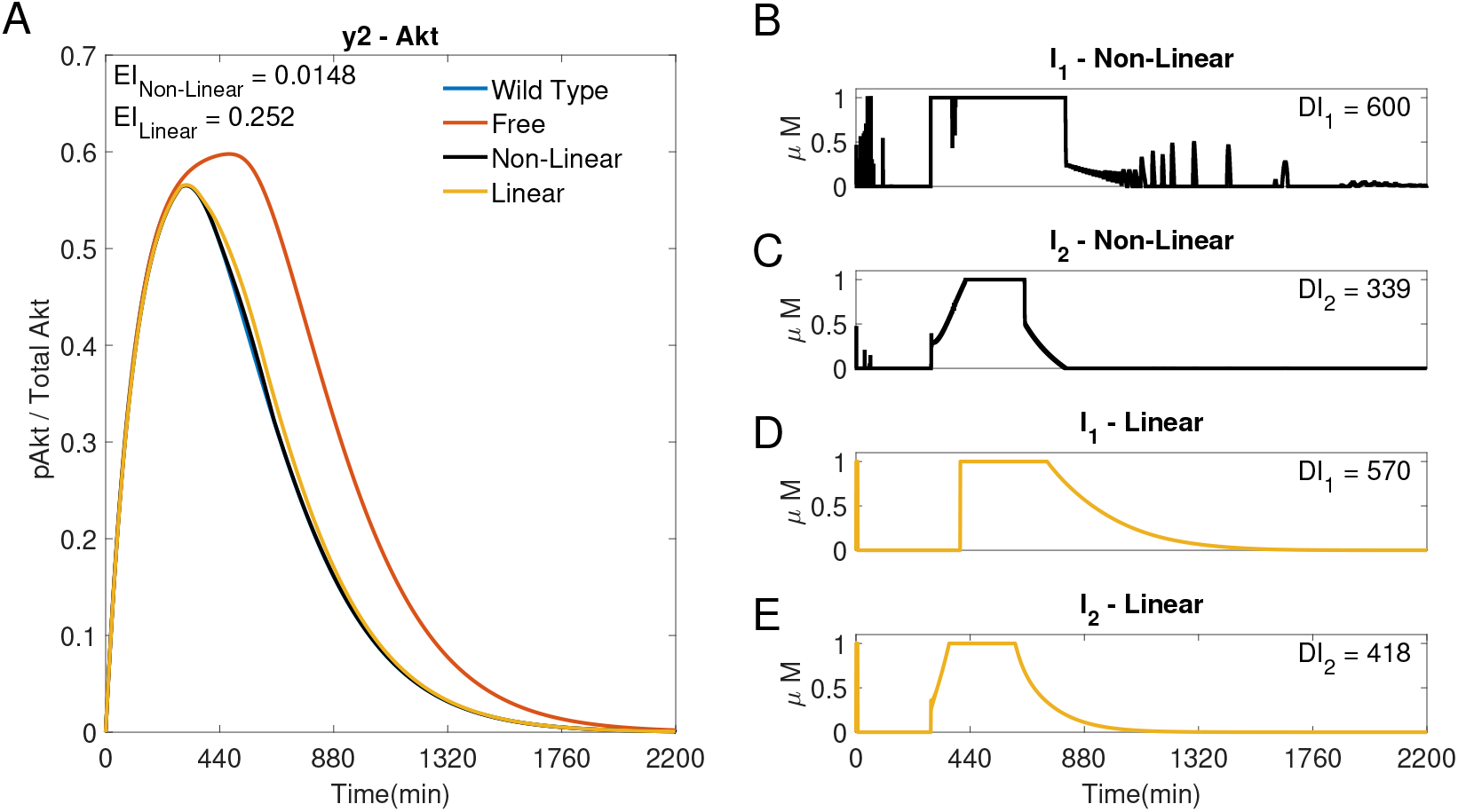
Two MISO MPC simulations using *I*_1_ and *I*_2_ to control the concentration of y2 (Akt). (A) The response of Akt to a non-linear MPC controller~ and an adaptive linear MPC controller ~. (B, C) Inputs *I*_1_ and *I*_2_ used in the non-linear simulation. (D, E) Inputs *I*_1_ and *I*_2_ used in the linear simulation. Parameters: *T_s_* = 1, *N* = 10, *α* = 0, *β* = 0, *γ* = [1, 10^5^, –], *θ* = 0 and *η* =1.

Figure S4 compares a MISO response using adaptive linear MPC ~ to non-linear MPC ~. It can be seen that the non-linear MPC has a lower Error Index of *EI* = 0.0148 compared to the adaptive linear MPC’s *EI* = 0.2520. However, the non-linear MPC had a significantly higher run-time, as expected.

Non-linear MPC would limit the controller’s use *in vitro*, as it the time to process the measurements might be longer than the data acquisition sampling time. This would then suggest using a larger sampling time, possibly causing issues with the controller performance (see Figures 5 and 6). When using the adaptive linear MPC controller, each iteration of the algorithm is well within the sampling time and enables capturing key dynamics of the system even though some of the non-linear couplings between the states are lost.

## S8 MPC vs Proportional (P) Control Schemes

All feedback simulations have used an MPC controller. Figure S5 compares the performance of a linear adaptive MPC controller ~ to a Proportional controller ~. Due to the relatively slow changing outputs, a differential gain was not used. An integral gain is not used as the output concentration is almost always greater than the reference, therefore the integral error never resets to zero, leaving the inputs at a non-zero steady state, causing a high *DI_i_*. Therefore a Proportional (P) controller is used. The two gains (for each input) can be tuned such that the P controller’s initial reaction to the state error results in a relatively low *EI*, as shown in Figure S5 ~, however the response is sensitive to the choice of gains. It can be seen in Figure S5 B) and C) that the inputs are identical, as the controller does not know the dynamics of the plant.

**Figure S5:**
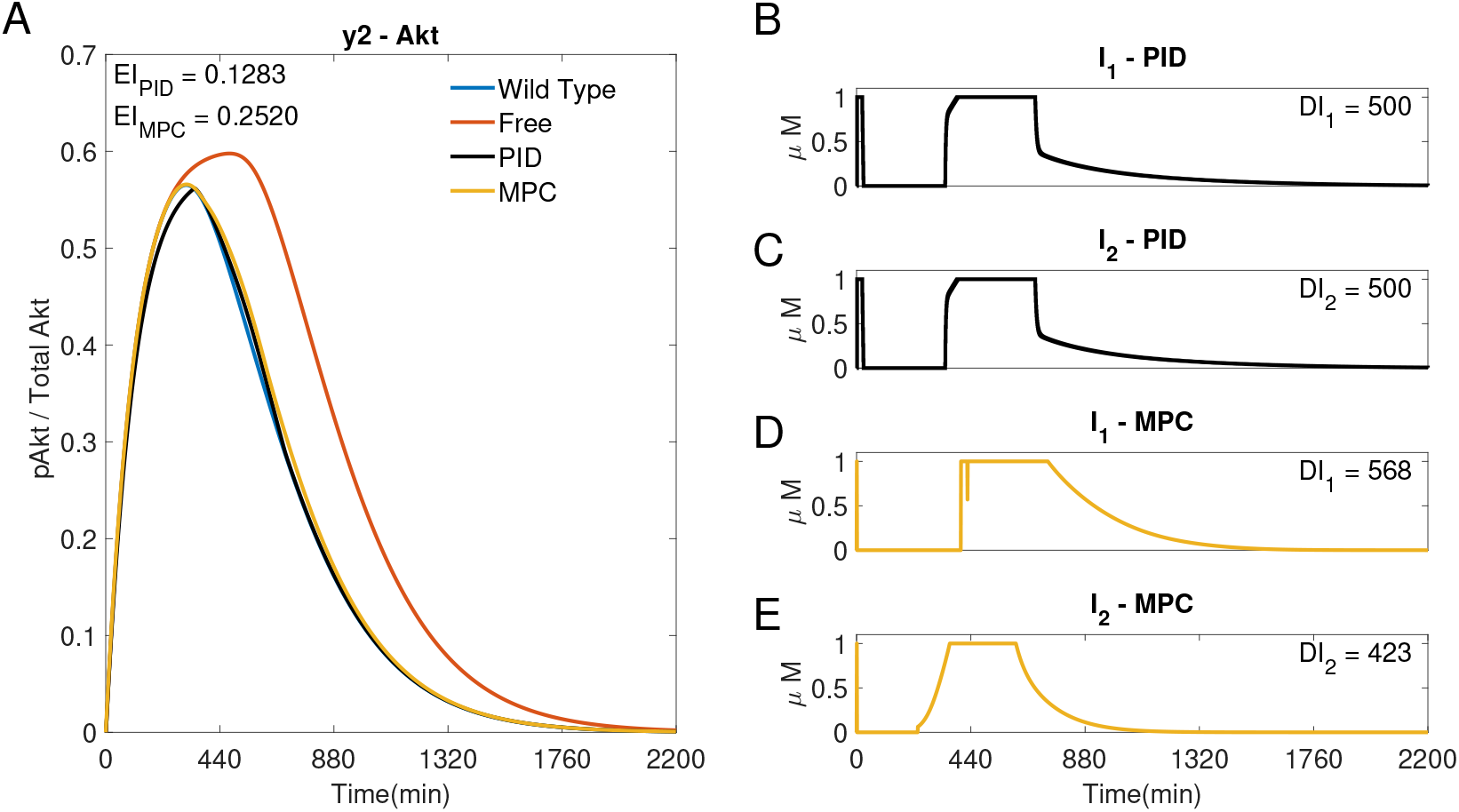
A comparison of a P~ and adaptive MPC ~ controller using *I*_1_ and *I*_2_ to control the concentration of y2(Akt). (A) The response of y2 (Akt) to a Proportional controller ~ and an adaptive linear MPC controller ~. (B, C) Inputs *I*_1_ and *I*_2_, respectively, used in the P simulation. =Proportional gains: *K*_*pI*_1__ = *K*_*pI*_2__ = 0.001. (D, E) Inputs *I*_1_ and *I*_2_ used in the MPC simulation. Parameters: *T_s_* = 1, *N* =10, *α* = 0, *β* = 0, *γ* = [1, 10^5^, –], *θ* = 0 and *η* = 1.

In Figure S5 the two controllers obtain a similar performance, but the P controller ~ offers no control on the inputs used. The P controller does not achieve robust control, as the low *EI* is as an effect of the finely tuned gains reacting to the initial error in the output, whereas the adaptive MPC controller ~ can be tailored for specific inputs’ choices.

## Notes

### Competing Interest Statement

The authors have declared no competing interest.

